# A plug-and-play ROI imaging module and deep-learning denoising framework extend three-photon microscopy to 1.7 mm depth

**DOI:** 10.64898/2026.01.02.697343

**Authors:** Jixiong Su, Shoupei Liu, Shasha Yang, Yanfeng Zhu, Xinyang Gu, Yaoguang Zhao, Chengyu Li, Min Zhang, Antao Chen, Huiyun Yu, Bo Li

**Author notes:** These authors contributed equally to this work (Jixiong Su, Shoupei Liu, Shasha Yang). Correspondence: Bo Li; Huiyun Yu.

## Abstract

Three-photon microscopy (3PM) has extended optical imaging beyond the reach of two-photon microscopy (2PM), but its practical depth remains far below theoretical predictions because of low photon efficiency and severe noise contamination. Here we present an integrated hardware and software solution that addresses these limitations. We developed a plug-and-play region-of-interest (ROI) imaging module that selectively excites neuron-occupied regions to improve power efficiency and reduce photothermal stress. The module incorporates frame-partitioned accumulative (FPA) imaging for high signal-to-noise ratio (SNR) structural acquisition, automated neuron segmentation, and motion-robust registration for stable recordings. Complementing this, OptiCal is a cascaded deep-learning framework that removes mixed noise including periodic, motion-induced, and random components. Together, these innovations enable high-fidelity imaging to 1.7 mm depth, extending the practical limit of 3PM by about 400 µm while maintaining low excitation power. Our results reveal reliable calcium activity and behavior-correlated dynamics in deep medial prefrontal cortex of awake mice.

## Introduction

Understanding how brain circuits operate requires real-time, high-resolution imaging deep within living tissue^1^. Two-photon microscopy (2PM) enables cortical imaging to depths of ~600 µm owing to reduced scattering and intrinsic optical sectioning^2–4^. Three-photon microscopy (3PM), which employs longer excitation wavelengths (1,300–1,700 nm) and higher-order nonlinearity, extends this range beyond 1 mm, reaching regions such as the hippocampus^5–7^, deep subregions of medial prefrontal cortex (mPFC)^8,9^, and even intact small organisms^10–12^. Despite these advances, achieving deeper functional imaging in scattering brain tissue remains a major challenge^13,14^.

Under ideal conditions, the signal-to-background ratio (SBR) serves as the criterion for the theoretical depth limit of 3PM, which can reach 3–4 mm in mouse brain vasculature^15^. In practice, however, two fundamental bottlenecks prevent the realization of this theoretical limit. First, deeper imaging requires higher excitation power, yet most excitation photons are scattered or absorbed in superficial tissue, generating heat rather than useful fluorescence^16–18^. This thermal effect restricts the maximum usable power and limits photon delivery to deeper regions^19–21^. Second, within this constrained power budget, the intrinsically small three-photon excitation cross-section yields weak fluorescence signals, causing rapid signal-to-noise ratio (SNR) degradation with increasing depth^22^.

Efforts to improve photon efficiency have taken two complementary directions. On the excitation side, adaptive illumination strategies selectively target regions of interest to reduce power demand^23^, but their implementation typically requires an additional synchronized laser source, limiting compatibility with existing three-photon systems and hindering widespread adoption. On the detection side, denoising algorithms originally developed for 2PM effectively suppress random fluctuations such as photon shot noise^24–27^, but they fail to account for the mixed-noise characteristics unique to three-photon imaging. At greater depths, where photon flux is extremely low, periodic structured noise from photomultiplier tubes (PMTs) and ambient illumination becomes a dominant artifact that further compromises signal fidelity^28,29^.

To overcome these limitations, we developed a plug-and-play region-of-interest (ROI) imaging module and a deep-learning-based denoising framework, OptiCal, to jointly enhance photon efficiency and signal fidelity in 3PM. The ROI module integrates frame-partitioned accumulative (FPA) imaging for high-SNR structural acquisition, automated neuron segmentation for rapid ROI generation, and an ROI imaging motion alignment algorithm (RIMA) for stable recordings in awake animals. Complementing these hardware advances, OptiCal employs a cascaded deep-learning architecture to remove mixed noise, including periodic, motion-induced, and random components, optimized for the low-photon-flux regime of three-photon imaging. Together, these innovations bridge the gap between theoretical and practical depth limits, extending the effective imaging depth of 3PM from ~1.3 mm to 1.7 mm. This represents an improvement of approximately 400 µm while maintaining low excitation power and high signal fidelity.

## Results

### Plug-and-play ROI imaging module for 3PM

To overcome the limited excitation efficiency that constrains deep-tissue 3PM, we developed a compact ROI imaging module capable of selectively exciting neuron-occupied regions to improve photon utilization and minimize photodamage **(Fig. 1a)**. This plug-and-play module integrates directly into existing three-photon systems **(Fig. 1b; Extended Data Fig. 1)**, combining a beam-gating hardware unit with dedicated control software supporting laser mode switching, automated segmentation, motion registration, deep-learning denoising, and signal extraction. The imaging workflow consists of three steps: (1) high-SNR structural acquisition using FPA imaging, (2) automated generation of ROI masks, and (3) real-time ROI-based functional imaging (**Fig. 1c**). A unified graphical user interface (GUI) integrates these functions and enables seamless control with a single click (**Fig. 1d**). With this software, we validated the precision of ROI pattern generation by scanning custom-defined shapes on fluorescent slides (**Fig. 1e**), and demonstrated *in vivo* that targeted excitation reliably captures neuronal and astrocytic activity (**Fig. 1f; Supplementary Video 1**). Together, these results establish a plug-and-play ROI imaging module that enables selective and efficient three-photon imaging on conventional microscopes, providing a hardware foundation for deep and automated ROI imaging.

**Fig. 1.**
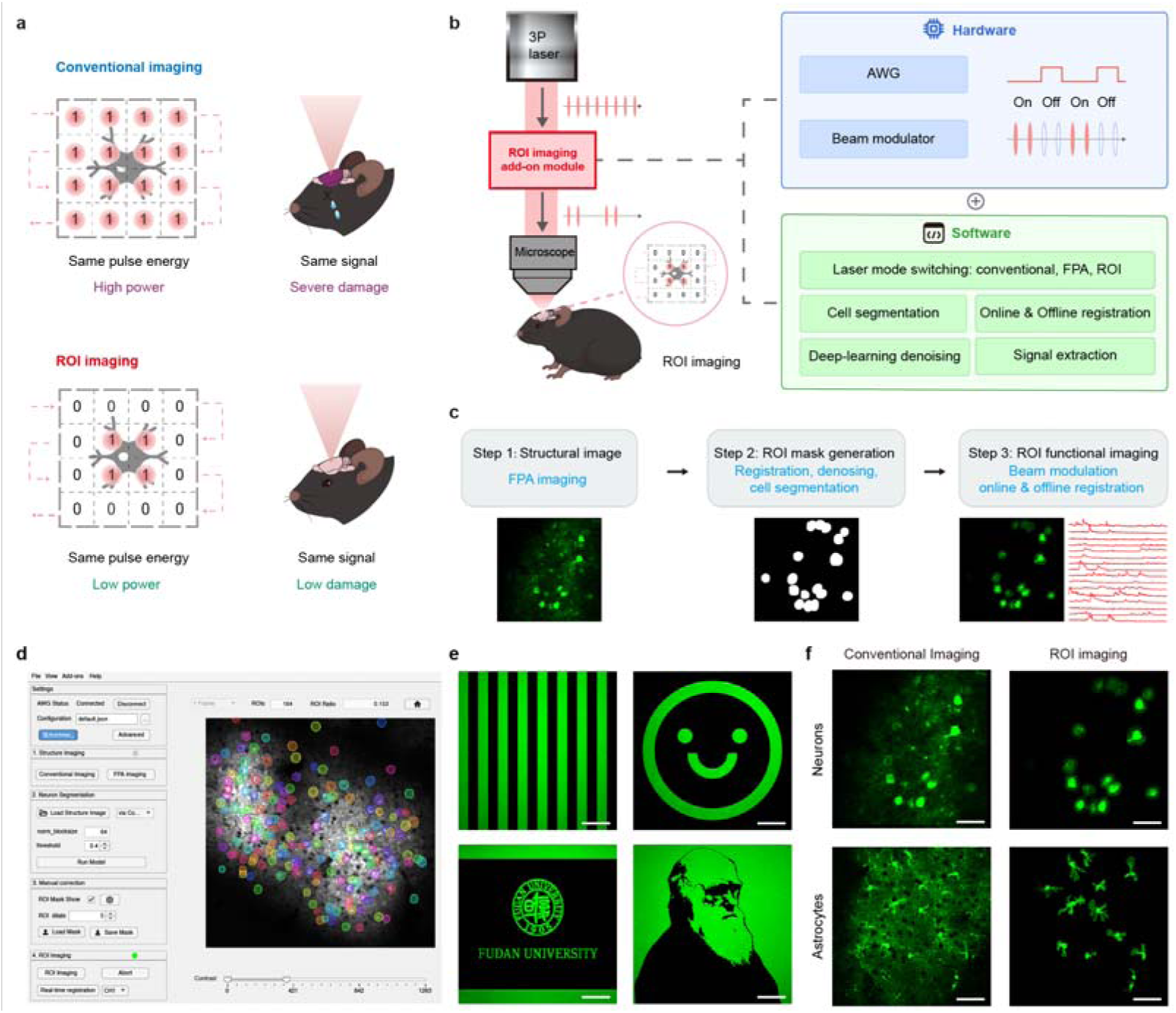
Design, workflow, and validation of the plug-and-play ROI imaging module for 3PM. **a**, Schematic comparison between conventional full-field imaging and ROI imaging. Conventional imaging (top) requires high peak power across the entire field-of-view (FOV), resulting in photobleaching and photodamage. ROI imaging (bottom) restricts excitation to cell-occupied regions, maintaining signal quality while substantially reducing the delivered average power. **b**, System schematic of the plug-and-play ROI imaging module. The module integrates a beam-gating unit, comprising an arbitrary waveform generator (AWG) and a beam modulator, with dedicated control software for synchronized operation. **c**, Overview of the ROI imaging workflow, including FPA structural imaging, automated ROI mask generation, and ROI-based functional imaging with online and offline registration. **d**, Control software interface for one-click switching between conventional and ROI imaging modes, with integrated controls for cell segmentation and motion registration. **e**, Validation of ROI pattern generation by scanning user-defined shapes on a uniform fluorescent slide. Scale bar, 50 μm. **f**, *In vivo* demonstration in awake mouse cortex. Conventional (left) versus ROI (right) imaging of GCaMP6s-labeled neurons (top, 539 μm depth) and astrocytes (bottom, 183 μm depth) illustrating targeted excitation and comparable fluorescence signals across both modes. Scale bar, 50 μm.

### Algorithmic innovations for deep, automated, and stable ROI imaging

Efficient ROI imaging in deep brain regions requires addressing three key technical challenges.

First, reliable identification of cellular targets at millimeter depths depends on structural images with sufficiently high SNR, which conventional full-field three-photon imaging fails to achieve due to limited photon yield and thermal constraints (**Fig. 2a, left**). To address this limitation, we developed FPA imaging, which restricts excitation to ~10% of the field of view per sub-frame, thereby increasing pulse energy tenfold without raising the average power. This higher pulse energy theoretically extends the effective imaging depth by ~2.3 attenuation lengths (~700 µm at 1,300 nm) and yields an approximately 10 dB improvement in SNR at equivalent depths while maintaining a much lower average power (**Fig. 2a-b**; **Extended Data Fig. 2; Supplementary Video 2**).

**Fig. 2.**
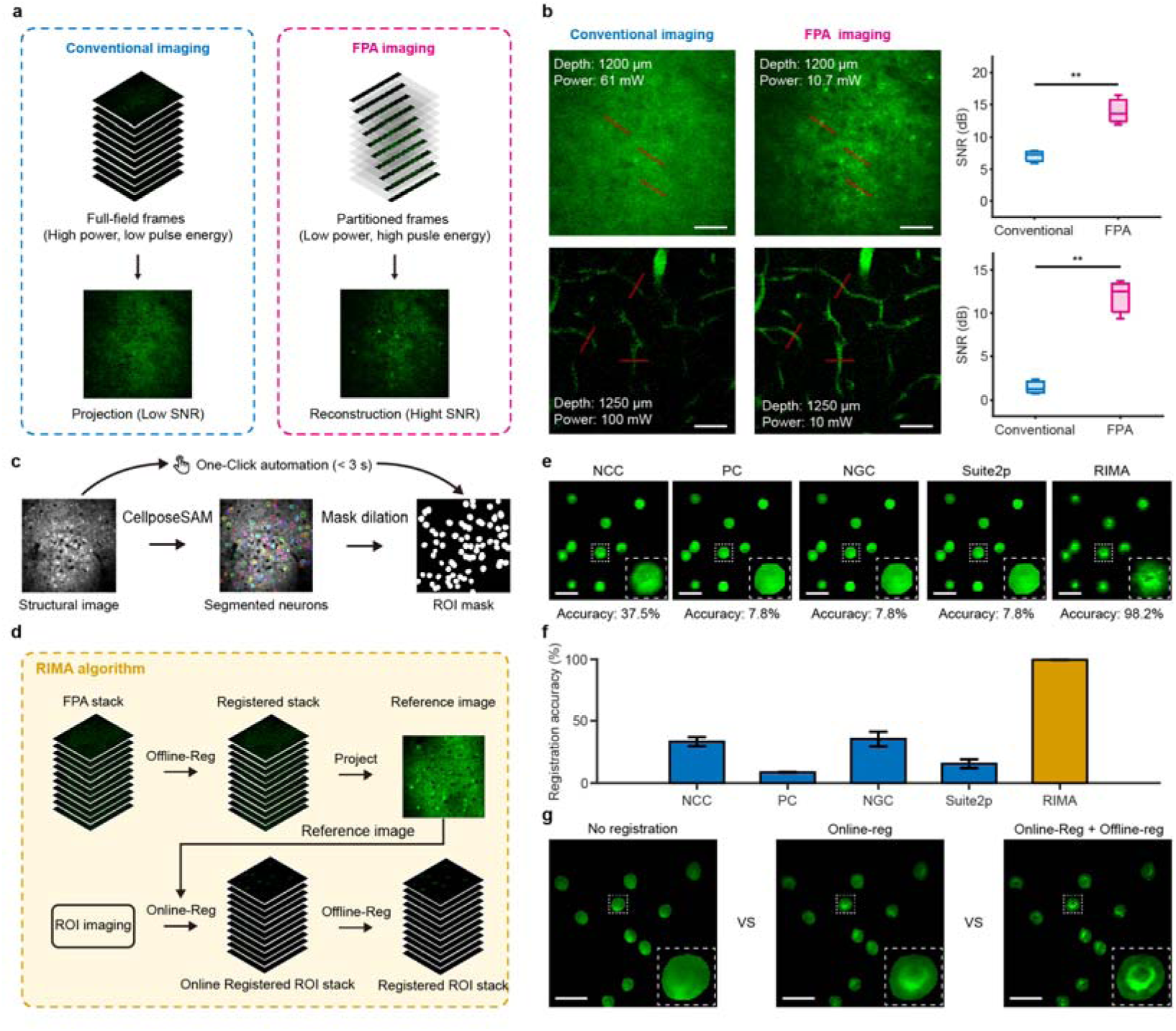
FPA imaging, automated segmentation, and RIMA enable robust deep-brain imaging. a, Schematic comparison of conventional and FPA imaging. In FPA imaging, each frame is subdivided into 10 sequential partitions, and a high-SNR full image is reconstructed by accumulating the 10 sub-frames. b, Comparison of SNR between conventional and FPA imaging in deep tissue. Left, representative maximum□intensity projection images of neurons (top, 1,200 µm; projection of 100 frames) and cerebrovasculature (bottom, 1,250 µm; projection of 10 frames) in the mouse brain with the two methods. Right, SNR quantification from the line profiles indicated in red (*n* = 3, \*\**p* < 0.01, two-tailed paired t-test). Scale bar, 50 μm. c, Automated workflow for one-click ROI mask generation. A high-SNR structural image is reconstructed from FPA imaging and segmented by CellposeSAM, the resulting mask is dilated to accommodate minor tissue displacements, and the binary ROI mask is generated for functional imaging. d, Workflow of RIMA. A motion-corrected reference is first created from an offline-registered FPA stack. This reference is then used to guide both online registration during ROI functional imaging and offline alignment of the recorded data. e, Comparison of RIMA and standard registration algorithms on simulated datasets with known motion. NCC: normalized cross-correlation, PC: phase correlation, NGC: normalized gradient correlation. Insets show magnified views of a single neuron. Scale bar, 50 μm. f, Quantification of registration accuracy across 50 datasets, showing RIMA’s superior performance. g, Comparison of imaging stability with no registration, online registration only, and combined online + offline registration. Insets show magnified views of a single neuron. The full RIMA pipeline maintains a stable field of view, which is essential for accurate signal extraction. Scale bar, 50 μm.

Second, in awake deep-brain recordings, the limited experimental time window renders manual ROI selection impractical. To automate this step, we implemented a rapid image-processing pipeline that reconstructs FPA-based structural images, performs neuron segmentation using CellposeSAM^30^, and generates ROI masks through ~5-µm dilation to accommodate minor physiological motion during imaging (**Fig. 2c**). The entire process completes within 3 s on our system, thereby substantially reducing preparation time for functional imaging in awake mice.

Third, brain motion during awake imaging introduces displacements in ROI-masked data that conventional registration algorithms cannot correct, as they assume all visible structures move coherently, a condition violated in ROI imaging, where the illumination pattern remains fixed while neurons drift. To overcome this mismatch, we developed RIMA, which aligns each ROI-masked frame to a motion-corrected, high-SNR FPA reference for motion stabilization (**Fig. 2d**). Benchmarking on simulated datasets with known displacements demonstrated 99.5% ± 0.3% alignment accuracy (*n* = 50), compared with < 40% for standard algorithms^31–34^ (**Fig. 2e,f; Extended Data Fig. 3; Supplementary Video 3**). The online implementation of RIMA ensures consistent ROI coverage throughout prolonged recordings (**Extended Data Fig. 4; Supplementary Video 4**), and, when combined with offline registration, the full RIMA pipeline maintains maximal spatial stability and signal fidelity across entire imaging sessions (**Fig. 2g**).

Together, these algorithmic advances, including FPA imaging for deep structural acquisition, automated segmentation for rapid ROI generation, and RIMA for motion-robust registration, collectively enable deep, automated, and stable ROI imaging at depths where conventional full-field 3PM fails.

### ROI imaging improves deep and large-FOV SNR while reducing phototoxicity

Having established the ROI imaging module, we next characterized its ability to improve signal quality for both deep and large-FOV imaging, while reducing phototoxicity in 3PM.

First, in deep-layer imaging, ROI excitation substantially improves signal quality while reducing power demand. In the motor cortex at 1,370 μm depth, conventional full-field imaging required 60 mW excitation power yet still produced low-SNR signals with indistinct calcium transients (**Fig. 3a**), a depth that typically marks the practical limit of conventional three-photon calcium imaging. In contrast, ROI imaging at only 12 mW, a fivefold reduction in power, produced well-resolved activity patterns from the same neuronal population (**Fig. 3b**) with markedly improved SNR (12.28 ± 1.36 dB versus 3.77 ± 0.91 dB; *n* = 10, *p* < 0.001, two-tailed paired t-test; **Fig. 3c**) and increased spike detection^35^ (39.70 ± 5.21 versus 0 ± 0; *n* = 10, *p* < 0.001, two-tailed paired t-test; **Fig. 3d**). These results indicate that selectively exciting neuron-occupied regions substantially enhances signal fidelity and photon efficiency in deep tissue, suggesting a potential route to surpassing the current depth limit of three-photon imaging.

**Fig. 3.**
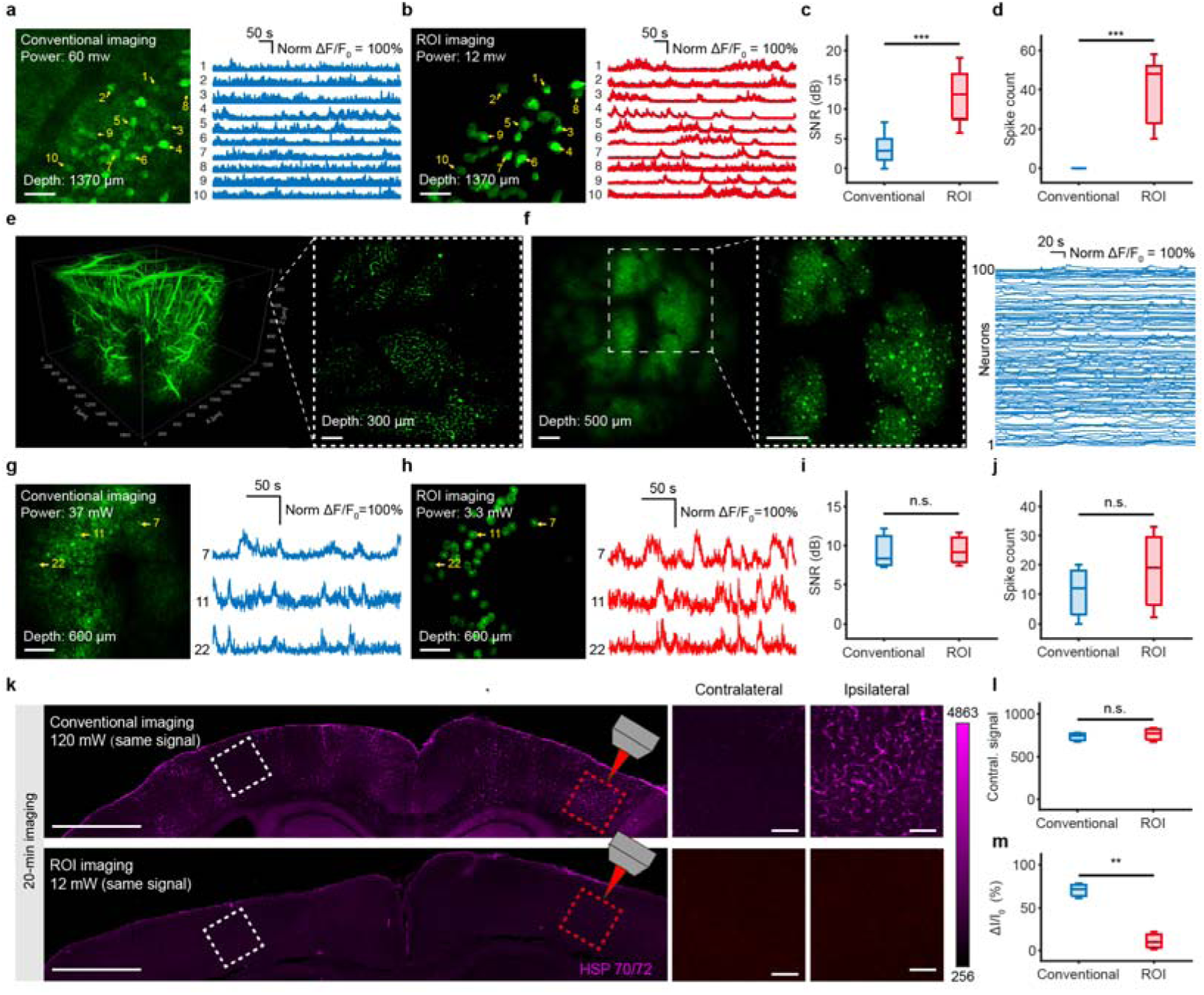
ROI imaging enhances signal fidelity, expands imaging scale, and reduces phototoxicity. **a**, Deep-layer neuronal imaging at 1370 μm under conventional imaging (60 mW). Left: structural image. Right: ΔF/F_0_ traces from 10 neurons. **b**, ROI imaging of the same neuronal population in **a** at 12 mW. **c**, Comparison of SNR for the ΔF/F_0_ traces in **a**,**b** (*n* = 10 neurons). **d**, Comparison of the number of inferred spikes for the ΔF/F_0_ traces in **a**,**b** (*n* = 10 neurons). **e,** Large-FOV three-photon structural imaging of mouse cerebrovasculature. Left, 3D reconstruction of a 1,970 × 1,970 × 1,200 µm^3^ volume (depth range: 0–1,200 μm). Right, representative cross-section at 300 µm depth. Scale bar, 200 µm. **f**, Large-area functional imaging of cortical neurons at 500 µm depth. Left, full FOV of 1,970 × 1,970 µm^2^. Middle, faster functional imaging with a zoomed-in FOV of 985 × 985 µm^2^ at 3.6 Hz. Right, ΔF/F_0_ traces from 100 neurons. Scale bar, 200 µm. **g**, Conventional full-field functional imaging of three representative neurons at 37 mW. Left, structural image. Right, corresponding ΔF/F_0_ traces from 3 neurons. Scale bar, 100 µm. **h**, ROI imaging of the same neurons as in **g** at 3.3 mW. Scale bar, 100 µm. **i**, Comparison of SNR for the ΔF/F_0_ traces in **g**,**h** (*n* = 3 neurons). **j**, Comparison of the number of inferred spikes for the ΔF/F_0_ traces in **g**,**h** (*n* = 3 neurons). **k**, HSP70/72 immunofluorescence after 20 min of continuous imaging at 1,000 μm. Images compare the effects of conventional (120 mW) and ROI (12 mW) imaging on the ipsilateral cortex. Left, coronal brain sections; Middle and right, magnified views of boxed regions. Scale bar in the left panel, 1 mm. Scale bar in the right panel, 100 μm. **l**, Quantification of mean fluorescence intensity (I_0_) in the contralateral hemisphere (*n* = 3 mice per group). **m**, Quantification of the relative intensity change (ΔI/I_0_) in the ipsilateral hemisphere (*n* = 3 mice per group). Statistical significance was assessed using two-tailed paired t-test in **c,d,i,j** and two-tailed independent t-test in **l,m**; \*\**p* < 0.01, \*\*\**p* < 0.001, n.s., not significant.

Second, for large-FOV imaging, ROI excitation similarly improves SNR while minimizing the power burden inherent to wide-area sampling. Wide-field 3PM typically requires high excitation power to capture sparse or distributed neuronal populations, increasing the risk of photodamage^36^. To address this, we integrated the ROI imaging module into a custom large-FOV three-photon microscope, achieving a 1,970 × 1,970 μm^2^ imaging area using a 16× objective (**Extended Data Fig. 6**). This configuration enabled volumetric reconstruction of cerebrovasculature down to 1,200 μm depth and wide-area functional imaging of hundreds of neurons (**Fig. 3e-f; Extended Data Fig. 7a-e; Supplementary Video 5**). Incorporating ROI imaging dramatically reduced the power requirement. Conventional scanning demanded 37 mW to capture robust calcium activity, whereas ROI imaging achieved comparable SNR at only 3.3 mW, a nearly tenfold reduction (**Fig. 3g-j**).

Finally, the reduced excitation power of ROI imaging directly translates into lower phototoxic stress. Immunofluorescence staining of HSP70/72 after 20 min of continuous imaging at 1,000 μm revealed strong expression following conventional imaging at 120 mW, indicating substantial photothermal damage. In contrast, ROI imaging at 12 mW produced minimal HSP70/72 expression, comparable to that in the contralateral, non-imaged hemisphere (**Fig. 3k-m**).

Altogether, these results demonstrate that ROI imaging enables high-fidelity functional recordings across both deep and large imaging regions, achieving an order-of-magnitude reduction in power consumption while effectively minimizing phototoxicity during extended *in vivo* imaging sessions. These findings establish the ROI imaging module as a key component of our deep three-photon imaging framework.

### OptiCal: a cascaded deep-learning model for mixed-noise removal in three-photon imaging

While ROI imaging maximizes excitation efficiency by directing photons only to neuron-occupied regions, efficient use of the emitted photons is equally critical for preserving overall signal fidelity. Existing denoising algorithms have been primarily developed for 2PM^24–27^, where noise is dominated by random fluctuations, particularly photon shot noise and background fluorescence. In contrast, three-photon imaging exhibits fundamentally different noise characteristics. Because of its smaller excitation cross section^37–39^, three-photon imaging operates under substantially lower photon flux, while its higher-order nonlinearity inherently suppresses background fluorescence^14,21^. Under these photon-limited conditions, small periodic or structured noise originating from photomultiplier tubes, ambient light leakage, or display flicker becomes a dominant source of signal degradation, particularly at greater imaging depths (**Fig. 4a**). Consequently, algorithms optimized for two-photon data, which assume predominantly random noise distributions, fail to generalize to the three-photon regime, leaving deep functional imaging severely noise-limited.

**Fig. 4.**
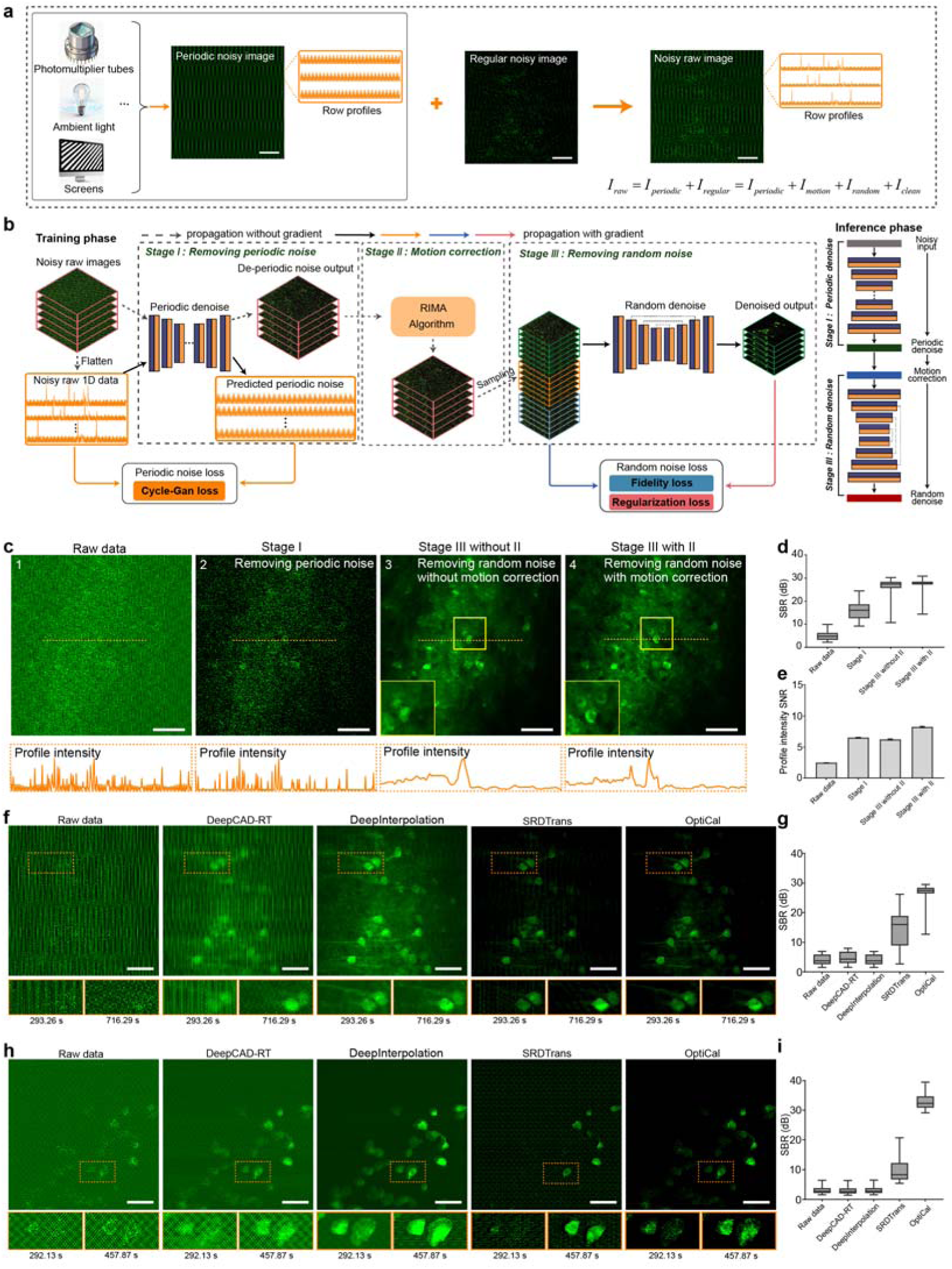
Principle of OptiCal and performance evaluation. **a**, Composition of raw noisy images (*I_raw_*) acquired by multiphoton microscopy, comprising periodic noise (*I_periodic_*), motion artifacts (*I_motion_*), random noise (*I_random_*), and clean fluorescent signals (*I_clean_*). **b**, Three-stage architecture of OptiCal and schematic of its training and inference workflow. **c**, Representative outputs from successive OptiCal stages for a three-photon image. Top (from left to right): raw data, periodic noise-removed image (Stage I), random noise-suppressed image without (Stage III without II) and with motion correction (Stage III with II). Bottom, corresponding intensity profiles along the orange dashed lines. **d**, Statistical comparisons of SBR across OptiCal stages (*n* = 31 regions of interest). **e**, Statistical comparisons of line-profile SNR across OptiCal stages (*n* = 512 rows). **f**, Representative three-photon images of GCaMP8s-labeled neurons processed by DeepCAD-RT, DeepInterpolation, SRDTrans and OptiCal. The bottom panels show magnified regions (orange boxes). **g**, Quantitative comparisons of SBR among denoising algorithms in **f** (*n* = 29 regions of interest). **h**, Representative ROI three-photon images of GCaMP6s-labeled neurons by DeepCAD-RT, DeepInterpolation, SRDTrans and OptiCal. The bottom panels show magnified regions (orange boxes). **i,** Quantitative comparisons of SBR among denoising algorithms in **h** (*n* = 32 regions of interest). Scale bar, 50 μm in **a**, **c**, **f**, **h**. Data are presented as mean ± SEM.

To address this challenge, we developed OptiCal, a cascaded deep-learning framework specifically optimized for three-photon imaging. OptiCal integrates three complementary stages to remove mixed noise sources characteristic of deep-tissue recordings (**Fig. 4b**). The first stage employs an unsupervised Cycle Generative Adversarial Network (CycleGAN) to eliminate periodic structured noise^40^. The second stage applies the RIMA algorithm to correct motion-induced artifacts. The final stage can incorporate existing denoising architectures. In this work, we reimplemented the transformer-based SRDTrans^27^ model and retrained it in an unsupervised manner on three-photon imaging data to suppress residual random noise and preserve fine structural details, yielding clean and high-fidelity calcium traces suitable for quantitative analysis.

To evaluate OptiCal’s performance, we performed *in vivo* three-photon calcium imaging of GCaMP6s-expressing neurons in the mouse somatosensory cortex at a depth of approximately 600 μm. The raw single-frame image was heavily contaminated by periodic and random noise, obscuring neuronal contours and introducing strong background fluctuations (**Fig. 4c1**). Sequential application of OptiCal progressively removed these artifacts: Stage 1 eliminated periodic structured noise **(Fig. 4c2**), Stage 2 corrected motion artifacts through RIMA integration, and Stage 3 suppressed residual random noise (**Fig. 4c3,4**). The final reconstruction revealed well-defined neuronal structures with smooth intensity profiles, demonstrating a marked improvement in structural clarity (**Supplementary Video 6**). Quantitative assessment of SBR and line-profile SNR across stages (**Fig. 4d,e**) confirmed significant enhancement in image quality.

To benchmark OptiCal against existing denoising approaches, we compared its performance with widely used algorithms originally developed for two-photon imaging, including DeepCAD-RT^26^, DeepInterpolation^24^, SRDTrans^27^. *In vivo* recordings of GCaMP8s-labeled neurons in the somatosensory cortex at a depth of 1,350 μm were processed using each method (**Fig. 4f**). While conventional algorithms effectively reduced random noise, they failed to remove periodic interference, leaving residual stripe artifacts and background fluorescence. In contrast, OptiCal simultaneously suppressed both structured and random noise, yielding clearer neuronal boundaries and artifact-free images. Quantitative analysis showed that OptiCal achieved >10 dB higher SBR compared with all other models (**Fig. 4g**).

We next evaluated the generalization of OptiCal under ROI imaging conditions, where mask-based excitation introduces additional structured and motion-related noise. Consistent with full-field results, OptiCal robustly suppressed periodic interference and random noise while maintaining stable signal intensity and neuronal morphology (**Fig. 4h; Supplementary Video 7**). Integration with RIMA ensured precise spatial alignment and a uniform background. Statistical comparison demonstrated an improvement exceeding 20 dB in fluorescence SBR relative to the next-best algorithm (**Fig. 4i**).

Together, these results establish OptiCal as a robust and generalizable framework for mixed-noise suppression in 3PM, enabling high-SNR reconstruction across both conventional and ROI imaging modes.

### High-SNR structural and functional imaging at 1.7 mm depth

With the ROI imaging module and OptiCal denoising framework established, we next evaluated the capability of our system for deep three-photon structural and functional imaging in the mouse mPFC. We injected pAAV-hSyn-GCaMP6s into the mPFC of C57BL/6J mice and performed *in vivo* imaging seven weeks later, when expression had stabilized (**Fig. 5a**).

**Fig. 5.**
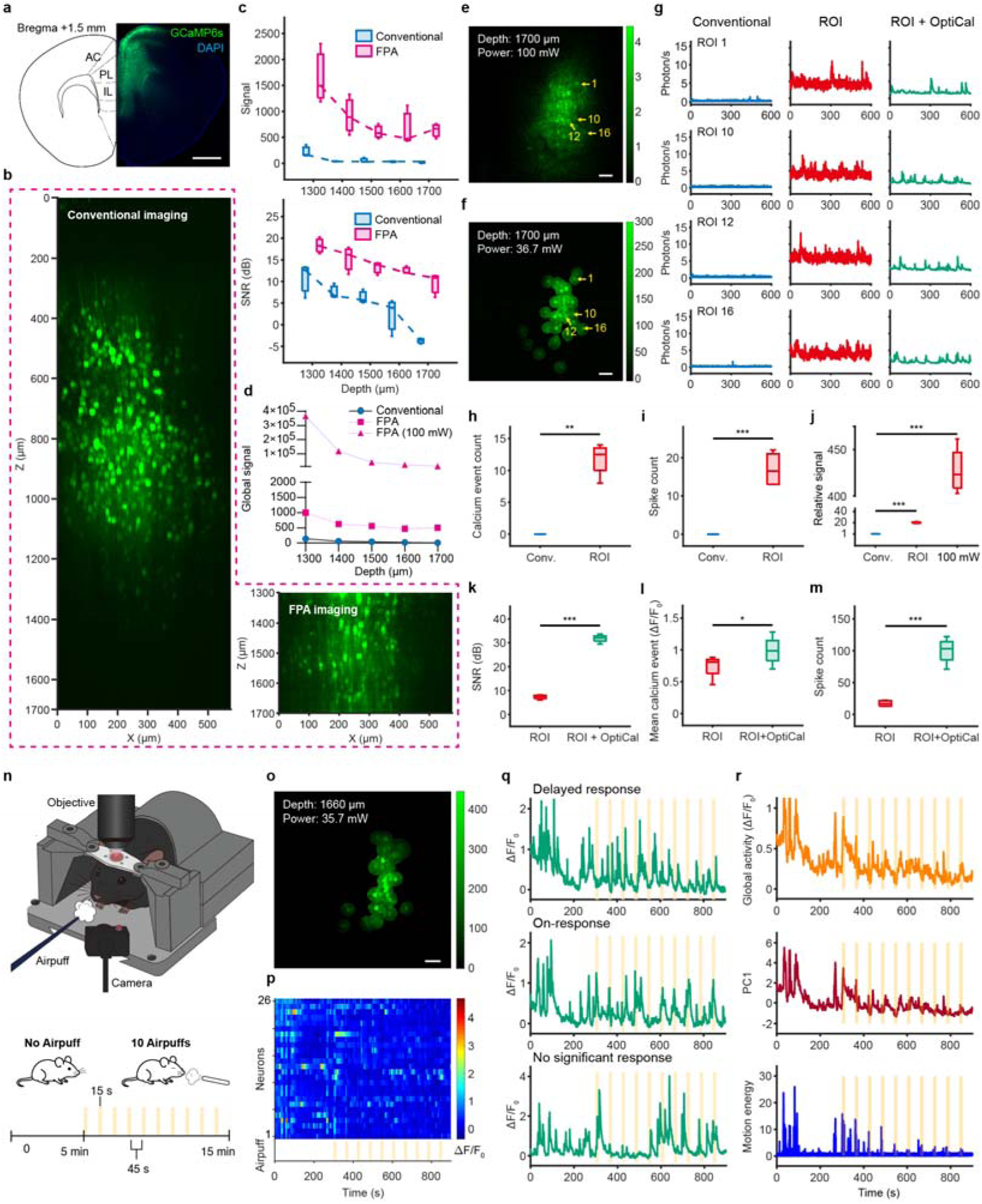
Deep mPFC neuron calcium imaging and functional analysis at 1,700 μm depth. **a**, Left: Schematic of mouse mPFC at Bregma +1.5 mm, showing anterior cingulate (AC), prelimbic (PL), and infralimbic (IL) cortices. Right: Representative histological image showing GCaMP6s expression (green) and DAPI counterstaining (blue). Scale bar, 1 mm. **b**, Maximum-intensity projection of volumes acquired with conventional (left) and FPA (right) imaging from 0-1,700 µm. FPA imaging maintained signal quality at reduced power (14-35 mW) compared with 100 mW for conventional imaging. **c**, Fluorescence signal (top) and SNR (bottom) across depths (1,300–1,700 µm) for the two imaging modes. **d**, Global signal intensity for conventional imaging, low-power FPA imaging, and theoretical FPA intensity extrapolated to 100 mW. **e,f,** Representative images of neurons at 1,700 µm depth acquired with conventional (100 mW) and ROI imaging. Scale bar, 50 µm. **g**, Calcium traces from four neurons indicated in **e** and **f**, comparing conventional, ROI, and ROI + OptiCal processing. **h,** Comparison of the number of detected calcium events between conventional and ROI imaging (*n* = 4 neurons). **i,** Comparison of the number of inferred spikes between conventional imaging and ROI imaging (*n* = 4 neurons). **j**, Relative fluorescence signal strength normalized to conventional imaging (set as 1). ROI: ROI imaging at 36.7 mW; 100 mW: ROI imaging power-normalized to 100 mW to match conventional imaging power (*n* = 4 neurons). **k**, Comparison of SNR for ROI traces before and after OptiCal denoising (*n* = 4 neurons). **l**, Comparison of calcium transient event amplitude for ROI traces before and after OptiCal denoising (*n* = 4 neurons). **m**, Comparison of the number of inferred spikes for ROI traces before and after OptiCal denoising (*n* = 4 neurons). **n**, Experimental setup for awake imaging (top) and air-puff stimulation paradigm (bottom). **o**, Representative FOV at 1,660 µm depth. Scale bar, 50 µm. **p**, Heatmap of calcium activity during the stimulation task. Yellow bars indicate air-puff stimuli. **q**, Example traces of three neurons showing delayed, on-response, and non-responsive activity. **r**, Population dynamics and behavior. Traces show the global average calcium activity, the first principal component (PC1), and motion energy extracted from behavioral data. Statistical significance was assessed using two-tailed paired t-test in **h-m**; \**p* < 0.05, \*\*\**p* < 0.001.

We first quantified the achievable imaging depth using conventional imaging and FPA imging. With conventional imaging, neuronal structures were clearly resolved up to approximately 1,300 μm (**Fig. 5b, left**), consistent with the practical depth limit commonly reported for 3PM. Beyond this depth, even at the maximum safe excitation power of 100 mW, fluorescence signals deteriorated rapidly, resulting in near-complete loss of structural visibility and sharp declines in signal intensity and SNR (**Fig. 5b, left; Fig. 5c**). In contrast, FPA imaging, operating at substantially lower powers (14–37 mW across the 1,300–1,700 μm range), enabled clear visualization of neuronal structures down to 1,700 µm (**Fig. 5b, right**). At matched depths, FPA imaging produced signal intensities 7.42-23.24 times higher than conventional imaging and maintained SNR values above 8 dB across the entire depth range, extending the effective imaging depth by ~400 µm beyond the conventional limit (**Fig. 5b, right; Fig. 5c, below; Extended Data Fig. 7a**). Given the cubic dependence of three-photon fluorescence on excitation power, extrapolation suggests that applying 100 mW within the FPA paradigm could yield up to ~1,000-fold stronger signals than conventional imaging at equivalent depths (**Fig. 5d**), highlighting its potential for even deeper imaging under optimized power conditions.

We next assessed calcium imaging performance at 1,700 μm by comparing conventional full-field scanning with our ROI-based imaging pipeline (**Fig. 5e,f**). Under conventional scanning, even at 100 mW excitation power, calcium signals from individual neurons were extremely weak (**Fig. 5g, left**). The average fluorescence intensity was only 0.21 ± 0.02 photons/s (**Fig. 5j**), rendering calcium transients nearly undetectable and precluding reliable spike inference (**Fig. 5h-i**). In contrast, ROI imaging of the same neurons, using only 36.7 mW (~one-third of the conventional power), produced substantially stronger signals (**Fig. 5g, middle)**, with an average fluorescence intensity of 4.65 ± 0.48 photons/s, representing a 21.62-fold increase (**Fig. 5j**). When normalized to the same power (100 mW), ROI imaging is projected to yield a 437.38-fold enhancement over conventional full-field imaging (**Fig. 5j**). These improvements enabled robust detection of 11.25 ± 1.38 calcium events per trial (**Fig. 5h**) and reliable inference of 17 ± 2.34 spikes per trial (**Fig. 5i**), revealing rich neuronal activity patterns inaccessible to conventional imaging.

Application of the OptiCal denoising further enhanced signal quality (**Extended Data Fig. 7b; Supplementary Video 8**). Calcium traces became cleaner and more interpretable (**Fig. 5g, right**), with SNR improving from 7.29 ± 0.45 dB to 31.79 ± 0.86 dB (16.79-fold enhancement; *n* = 4, *p* < 0.001, two-tailed paired t-test; **Fig. 5k**). Denoising also increased the amplitude of calcium transients (ΔF/F_0_: 0.7381 ± 0.10 to 0.99 ± 0.12; *n* = 4, *p* < 0.001, two-tailed paired t-test; **Fig. 5l**), facilitating more reliable event detection. Correspondingly, the number of inferred spikes rose dramatically, from 17 ± 2.34 to 99.75 ± 10.15 spikes per trial (*n* = 4, *p* < 0.001, two-tailed paired t-test; **Fig. 5m**), underscoring the combined benefits of improved SNR and transient fidelity.

Collectively, these results show that the integration of FPA imaging, ROI-based excitation, and OptiCal denoising enables reliable, high-SNR structural and functional imaging at depths up to 1.7 mm in the awake mouse brain. This represents an approximately 400 µm extension beyond the conventional practical depth limit of 3PM, achieved at significantly lower excitation power.

### Population-level behavioral encoding in deep mPFC

Having established stable deep-layer recording, we next examined whether our imaging pipeline could capture population-level neural dynamics related to behavior. The infralimbic (IL) subdivision of the mPFC, located beyond 1,500 μm, was targeted for *in vivo* imaging in awake, head-fixed mice (**Fig. 5n**). Previous optical studies of this region have been restricted to ~1,100 μm or required invasive removal of overlying cortex^8,41^. Using our ROI-based three-photon system combined with OptiCal denoising, we achieved stable calcium imaging in this deep region. Each session included a 5-min baseline followed by 10 min of intermittent air-puff stimulation (15 s stimulation, 45 s inter-trial interval; **Fig. 5n**). The entire 15-min recording was acquired at an average power of only 35.7 mW, demonstrating safe and sustained imaging under deep-tissue conditions (**Fig. 5o**).

The resulting calcium activity maps revealed heterogeneous responses, including both stimulus-driven and delayed activity patterns (**Fig. 5p-q**). Although detailed circuit mapping was beyond the scope of this study, the high data quality enabled extraction of population-level activity components. Principal component analysis (PCA) of ΔF/F_0_ signals showed that the first component (PC1) captured the dominant shared population variance and was highly correlated with the global average activity (r = 0.9863; **Fig. 5r**), reflecting strong synchrony across neurons. To assess behavioral relevance, we quantified movement energy from simultaneously acquired behavioral videos. Both the global ΔF/F_0_ trace and PC1 were significantly correlated with movement energy (r = 0.6564 and r = 0.6965, respectively; **Fig. 5r**), indicating that deep mPFC population activity is modulated by behavioral state.

Together, these findings demonstrate that our ROI-based three-photon and OptiCal-enhanced imaging framework provides the stability, sensitivity, and signal quality necessary to uncover population-level behavioral representations at depths approaching 1.7 mm in the intact mouse brain.

## Discussion

3PM has opened new opportunities for deep-tissue imaging^15^, yet its practical performance has remained far below theoretical expectations due to poor photon efficiency and severe noise contamination^21,22^. In this work, we present a plug-and-play ROI imaging module and a deep-learning denoising framework, OptiCal, that jointly overcome these long-standing barriers. Together, these tools extend the effective imaging depth of 3PM from the conventional limit of ~1.3 mm to 1.7 mm while maintaining low excitation power and high signal fidelity. Importantly, both components are compatible with existing microscope architectures and can be readily adopted for routine *in vivo* applications.

The concept of ROI imaging builds on our previously developed adaptive excitation source (AES) system^23^, which dynamically redistributes pulse energy to neuron-occupied regions to enhance excitation efficiency. While AES demonstrated the feasibility of region-selective three-photon excitation, several practical limitations remained. First, AES required replacement of the laser source rather than simple integration into existing microscopes. Second, its accessible wavelength range was restricted to 1,700 nm due to soliton self-frequency shift (SSFS) effects^42^, precluding efficient excitation of GFP-based fluorophores at 1,300 nm^22^. Third, the system focused primarily on excitation optimization and did not address key challenges such as deep structural image acquisition, automated neuron segmentation, or motion-robust registration. Finally, noise suppression and signal fidelity at depth were not optimized. The present work overcomes these limitations by introducing a modular hardware design and a cascaded deep-learning framework for comprehensive optimization of both excitation and detection.

Our approach is also complementary to optical correction techniques such as adaptive optics (AO)^43–45^. AO compensates for tissue-induced aberrations to restore focus and improve signal strength, whereas ROI imaging minimizes unnecessary excitation and photothermal load. The two strategies address distinct physical bottlenecks and are inherently synergistic. Since AO typically requires preliminary structural images to estimate aberrations, this dataset can directly serve as the structural reference for ROI mask generation, enhancing both correction accuracy and system efficiency. Integration of ROI imaging and AO may thus push the practical depth of 3PM even closer to its theoretical limit.

On the detection side, OptiCal introduces a new paradigm for denoising in multiphoton imaging. Unlike conventional algorithms that primarily target random fluctuations^24–27^, OptiCal performs sequential suppression of periodic, motion-induced, and random noise through a cascaded deep-learning framework. Although optimized for three-photon data, its unsupervised and modular design makes it broadly applicable to other low-photon-flux modalities, including label-free harmonic generation^46^, Raman scattering^47^, and low-power fluorescence endoscopy^48^.

While we demonstrated high-SNR structural and functional imaging at 1.7 mm depth using only 37 mW of excitation power, this depth does not represent a fundamental limit of our system. The maximum achievable depth in the current study was constrained by the 600 mW output of our laser source, which limited the available power after ROI modulation. In contrast, most commercial three-photon lasers deliver 1–4 W of average power, enabling effective ROI imaging with powers exceeding 100 mW. Considering the cubic dependence of three-photon fluorescence on excitation power^37,39^, such configurations could feasibly extend the imaging depth beyond 2 mm under safe operating conditions.

Finally, both hardware and software components of our system were developed with broad accessibility in mind. The ROI module can be inserted between the laser and the microscope using standard acousto- or electro-optic modulators already present in most commercial three-photon lasers. OptiCal is available as a standalone software package and can be applied directly to existing datasets without hardware modification. Together, these features establish a practical, scalable, and easily disseminated framework for deep, high-fidelity functional imaging in the intact mammalian brain.

## Methods

### Imaging system configuration

#### Conventional 3PM setup

Our system was built on a custom-designed upright microscope. Three-photon excitation was provided by a non-collinear optical parametric amplifier (I-OPA-TW-F, Light Conversion) pumped by a 40 W femtosecond laser (Carbide-CB3, Light Conversion). The system was tuned to an output wavelength of 1,300 nm with a 1 MHz repetition rate.

Beam scanning was performed by a galvanometer scanner (6210H, Novanta). Fluorescence and third-harmonic generation (THG) signals were collected in the epi-direction through the objective. The emission light was first reflected by a primary dichroic mirror (FF775-Di01, Semrock) and subsequently separated by a secondary dichroic mirror (FF458-Di02, Semrock). The fluorescence and THG signals were then passed through respective bandpass filters (Fluorescence: 520/60 nm, Semrock; THG: 447/60 nm, Semrock) before being detected by independent photomultiplier tubes (PMTs; H7422-40, Hamamatsu). The PMT output currents were converted to voltages by a transimpedance amplifier (C12419, Hamamatsu). The sample was mounted on a motorized three-axis stage (MP-285A, Sutter Instrument).

System control and image acquisition were managed by ScanImage (v2022.1.0, Vidrio Technologies) running in MATLAB (R2023b, MathWorks) on a workstation equipped with an Intel Core i9-13900KF CPU, 96 GB of RAM (5600 MHz), and an NVIDIA GeForce RTX 4090 graphics card (24 GB) running a 64-bit Windows operating system.

Imaging was performed with a 25× water-immersion objective (NA = 1.05, 2 mm working distance; XLPLN25XWMP2, Olympus), giving a FOV of 285 × 285 μm^2^ (zoom = 2). Images were acquired at 512 × 512 pixels and 3.56 Hz.

#### Large-FOV 3PM setup

For large-FOV imaging, we used an integrated femtosecond laser (CRONUS-3P, Light Conversion) delivering 1,300 nm pulses at 1 MHz repetition rate. The custom wide-field scanning module consisted of a high-speed galvanometer scanner (6215H, 5 mm aperture, Cambridge Technology) operating at 1.792 kHz for the X-axis (±7.5° range). The scanned beam was relayed through a scan lens (*f* = 110 mm) and tube lens (*f* = 200 mm). All other optical and detection components were identical to those in the conventional 3PM setup.

Two water-immersion objectives were used: a 25× objective (NA = 1.05, 2 mm working distance; XLPLN25XWMP2, Olympus) and a 16× objective (NA = 0.8, 3 mm working distance; CFI75 LWD 16X W, Nikon). The 25× objective provided a FOV of 1,152 × 1,152 μm^2^ (zoom = 1). The 16× objective provided a FOV of 1,970 × 1,970 μm^2^ (zoom = 1). Images were acquired at a resolution of 512 × 512 pixels, yielding frame rates of 1.80 Hz at zoom = 1 and 3.60 Hz at zoom = 2.

### FOV and resolution measurement

The FOV of our custom-built three-photon microscope was determined using a standard microscope calibration slide (100 μm division, 0-20 mm measurement range; AOSVI, Shenzhen, China).

Spatial resolution was measured as the point spread function (PSF) of 0.05–0.1 μm green fluorescent microspheres (Lumisphere, BaseLine Chrom Tech Research Centre, Tianjin, China). The microspheres were diluted in physiological saline from a 1% stock suspension. Following three-photon imaging, the lateral and axial intensity profiles of single microspheres were normalized and smoothed with a Gaussian filter. The resolution in the X, Y, and Z dimensions was then defined as the 1/e radius of the smoothed intensity profile (Extended Data Fig. 5c-d,f-g).

### Development of ROI imaging module

#### Hardware setup

The core of this module is an Arbitrary Waveform Generator (AWG; PXI-5421, National Instruments), housed in a PXIe chassis (PXIe-1062Q, National Instruments). The control computer interfaces with the PXI chassis via a PXIe-PCIe8361 remote controller, allowing custom scripts to directly command the AWG.

To ensure precise synchronization, the system was configured as follows. First, the clock input (CLK IN) of the AWG was connected to the 1-MHz master clock output of the laser system, synchronizing the AWG sample rate with the laser repetition rate for one-pulse-per-pixel imaging. Second, the programmable function interface (PFI0) of the AWG was connected to the “Frame Start Trigger” output of the ScanImage vDAQ system, ensuring that waveform generation began precisely at the start of each acquisition frame. Finally, the AWG channel 0 (CH0) output was connected to the analog modulation input of the laser’s integrated acousto-optic modulator (AOM). The AOM was configured for active-low logic, where a 0 V signal from the AWG activated the laser, and a 3.3 V signal suppressed the laser.

#### Beam modulation signal generation

To generate the laser modulation signal, 2D ROI masks containing neuron coordinates were converted into a 1D logical sequence (Extended Data Fig. 1b). This conversion algorithm accounted for the non-imaging turnaround time of the galvanometer scanner at the lateral boundaries of the field of view during bidirectional scanning. A calibrated delay was introduced after the frame trigger to synchronize the laser modulation with the physical position of the scanners. The laser was programmed to deactivate immediately after scanning the final pixel of the last ROI in a frame. This pixel-targeted excitation strategy minimized total optical energy deposition and reduced potential phototoxicity. The resulting modulation signals were streamed to the laser on a frame-by-frame basis.

#### Conventional registration algorithms

##### Normalized Cross-Correlation (NCC)^31^

This is a spatial domain method that calculates similarity between reference and target images. For displacement (*u, v*):

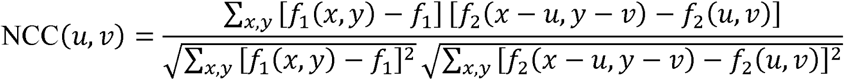

where *f*_1_ is the reference image, *f*_2_ is the target image, *f*_1_ is the reference image mean, and *f*_2_(*u, v*), is the target image window mean at the current displacement.

##### Phase Correlation (PC)^32^

This is a frequency domain method that is robust to brightness changes. It uses the Fourier transform shift property to determine displacement via the cross-power spectrum:

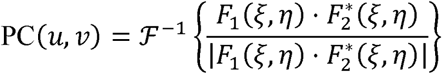

where *F*_1_ and *F*_2_ are the Fourier transforms of the two images, 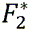 is the complex conjugate of *F*_2_, (*ξ*, *η*), represents frequency domain coordinates, and (*u*, *v*) represents the displacement. Peak location in the correlation plane determines the displacement.

##### Normalized Gradient Correlation (NGC)^33^

This method compares image gradient fields rather than pixel intensities, providing insensitivity to brightness changes. The method involves optional Gaussian smoothing to reduce noise, followed by gradient calculation *G_x_* and *G_y_* in x and y directions, which are combined into a complex gradient image, *G*(*x*, *y*) = *G_x_*(*x*, *y*) + *i · G_y_* (*x*, *y*). The normalized cross-correlation is then computed for the complex gradient images *G*_1_ and *G*_2_:

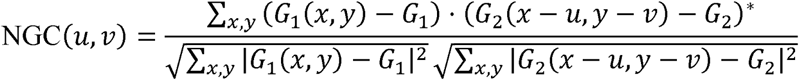

where *G*_1_ and *G*_2_ are the complex gradient image means, and * denotes complex conjugate.

##### Suite2p^34^

This is a widely used two-photon calcium imaging processing package. Its registration algorithm improves upon standard phase correlation by applying Gaussian filtering to remove high-frequency noise and using sub-pixel interpolation for enhanced accuracy. Instead of using a single frame as reference, Suite2p iteratively computes an average reference image from the aligned stack.

#### Registration algorithms for ROI imaging

Conventional registration algorithms show a significant drop in performance when processing the ROI-masked data. To address this, we developed RIMA, a registration algorithm specifically designed for ROI-masked data.

RIMA is a registration strategy based on the NGC algorithm. It utilizes a high-SNR, full-field reference image derived from a stack acquired via FPA imaging before the ROI imaging session.

This reference image is created using the following iterative refinement process. First, the single frame from the full-field stack with the highest mean correlation to all other frames is identified as the initial reference. Second, the entire stack is registered to this initial reference using the NGC algorithm. Third, the 20 frames exhibiting the highest correlation from this newly registered stack are selected and averaged to create an updated, higher-SNR reference. This two-step refinement process of registering the stack and averaging the top 20 frames is iterated 8 times. The average image resulting from the final iteration serves as the definitive full-field reference.

During the subsequent registration for ROI imaging data, each incoming ROI-masked frame is aligned to this full-field reference.

#### Quantification of registration accuracy

Registration accuracy is defined as the percentage of frames where the squared Euclidean distance between the predicted displacement and the true displacement is 10 pixels or less. All experiments are based on *n* = 50 calcium imaging stacks with applied displacements. The dataset used for benchmarking was generated by applying random displacements to real, conventionally acquired calcium imaging data to provide a ground truth for performance evaluation. The simulated ROI imaging data was then produced by applying a fixed ROI mask to these displaced image stacks.

#### Online registration for ROI imaging

Because the RIMA algorithm generates a full-field reference image containing data from the entire field of view, it can be used for both offline and online registration.

We implemented online registration functionality based on the RIMA algorithm by integrating it with ScanImage’s built-in registration framework. The registration process is executed at user-defined time intervals or when image displacement surpasses a specified threshold. The computed x-y offset is then fed back to the galvanometer scanners, which apply a corrective angular shift to the scan pattern. This galvo-based correction mechanism enables high-speed, low-latency motion compensation.

#### Automated segmentation to generate ROI masks

To automatically identify neuronal cell bodies for ROI imaging from structural images with high SNR, we employed a deep learning-based segmentation workflow.

This workflow is built on the Cellpose-SAM model^30^. The model combines the Cellpose framework with pre-trained Segment Anything model weights, featuring exceptional generalization capabilities that exceed inter-rater annotation consistency. We achieved favorable results by using the original model directly without additional training.

After acquiring three-photon image stacks, motion correction was performed on them. Subsequently, three types of projection maps were computed: average projection, maximum projection, and standard deviation projection. One of these projection maps was selected, and the image was input into the Cellpose-SAM model. Key segmentation parameters were optimized for our dataset: the normalization block size (norm_blocksize) was set to 64 to handle uneven illumination in images; the cell detection threshold (threshold) was set to 0.4 to balance detection sensitivity and false positive rate.

#### Software development

We developed custom GUI software using the App Designer in MATLAB (R2023b, MathWorks).

The software features bidirectional communication with ScanImage, allowing it to query system status and retrieve key parameters such as FOV, resolution, and frame rate. These parameters are utilized to calibrate the output of the AWG.

Based on user selection, the software can generate and output scanning waveforms for three distinct modes with a single click: conventional imaging, FPA imaging, and ROI imaging. In FPA mode, the software automatically stitches the acquired sub-frames in real-time to reconstruct a complete structural image for preview.

The software integrates the automated cell segmentation function described above, allowing users to invoke the Cellpose-SAM model and automatically load the resulting ROIs with one click. Furthermore, the software provides a complete suite of manual ROI tools, enabling users to add, modify, or delete ROIs directly. Once ROIs are finalized, the software automatically applies a ~5-µm dilation to each mask. This dilation accommodates minor physiological motion during imaging

Additionally, the software also implements the RIMA-based online registration, which can be activated with one click to enable motion correction during live imaging sessions.

### 3PM deep learning-based denoising

#### Network architecture of OptiCal

OptiCal adopts a three-stage cascaded architecture, which decomposes the denoising task of low SNR 3D imaging data into three sequential and independent subtasks: periodic noise elimination, motion blur correction, and random noise elimination. Each stage is optimized for a single type of noise, which not only improves training efficiency but also achieves comprehensive suppression of multiple noise types in the original data.

For the input low-SNR 3D imaging data (with dimensions *H* × *W* × *T*, where *H* = height, *W* = width, and *T* = number of frames), the first stage constructs a training dataset through preprocessing: the original 3D data undergoes frame-level row-wise unfolding preprocessing, where each 2D frame is unfolded pixel-by-pixel along rows into a 1D vector to generate a 1D original data subset. Meanwhile, pre-collected 1D periodic noise data matched to the imaging system is introduced. Together, these two components form an unpaired training dataset (Set A: 1D original data; Set B: 1D periodic noise data).

This unpaired dataset is fed into the CycleGAN network for training: On one hand, the 1D original data is first input to Generator G, which outputs predicted 1D periodic noise data. This predicted data is then fed into Generator F and Discriminator D_B_ to calculate the cycle-consistency loss and adversarial loss, respectively. On the other hand, the 1D periodic noise data is synchronously input to Generator F, which outputs predicted 1D original data. This predicted data is further fed into Generator G and Discriminator D_A_, with the same two types of losses calculated. After the training converges, the CycleGAN can realize bidirectional mapping between the 1D original data and the periodic noise. At this point, subtracting the predicted periodic noise output by Generator G from the original data yields the output data with periodic noise eliminated (De-periodic data).

To eliminate motion blur noise caused by sample movement and avoid its interference with subsequent random noise elimination, the second stage of OptiCal employs RIMA algorithm for conventional and ROI imaging, achieving inter-frame alignment correction for the De-periodic data.

The third stage focuses on random noise elimination for the corrected data, which only contains of effective signals with independently distributed random noise. Given the statistical property that independent random noise converges to its mean estimate through extensive training, this stage allows flexible selection of suitable high-performance denoising algorithms (e.g., DeepCAD, DeepCAD-RT, SRDTrans). Considering SRDTrans’ advantages in random noise suppression and computational efficiency for high-dimensional biological imaging data, this study selects SRDTrans as the denoising algorithm for the third stage.

#### Model training and inference

The training and inference of the OptiCal model are performed in a specific hardware and software environment: the hardware is a workstation equipped with an Intel Xeon Gold 6248R processor (@3.00 GHz) and an NVIDIA A5000 GPU; the software environment is built based on Python v3.10.13 and the PyTorch v2.1.0 framework.

In the periodic noise elimination stage, data preprocessing is first performed before training: both the original low-SNR 3D data and the pre-collected periodic noise data undergo 1D transformation.Spifically, each 2D frame in the 3D data is unfolded pixel-by-pixel along rows into a 1D vector with a dimension of 512 × 1. Finally, an unpaired training set containing 10,000 samples is constructed (Set A: 1D original data; Set B: 1D periodic noise data).

For training parameters, the Adam optimizer is used (β = 0.9, β = 0.999, weight decay coefficient = 1e-5), with a learning rate set to 1e-4, a training batch size of 10, and a total of 30 training epochs.

During the inference process after training, the original low-SNR 3D data to be processed is unfolded into 1D vectors in the same manner and input to the converged Generator G, yielding predicted periodic noise vectors. Subtracting these noise vectors from the original 1D data and reconstructing the result into 3D data completes the periodic noise elimination.

After periodic noise elimination and motion correction, the 3D data is used as input to train the SRDTrans model for the random noise elimination stage. The training parameters are set as follows: a learning rate of 1e-4, a training dataset containing 6000 3D data blocks (each data block has dimensions of 128 × 128 × 128 pixels), and a total of 30 training epochs.

The periodic noise elimination model of OptiCal exhibits system dependence: under the same imaging system, it only needs to be trained once, and subsequent data collected by the same system can directly call this model for inference. The random noise elimination model exhibits sample type dependence: for the same type of imaging samples, it only needs to be trained once, and data from similar samples can directly reuse this model.

#### SBR calculation

To quantitatively evaluate the denoising effect, the SBR of each image stack is calculated according to the following method: the average fluorescence intensity of the ROI is selected as the signal (*S*), and the average intensity of other unstructured background regions is selected as the random noise intensity (*B*). The SBR (dB) of the image is then calculated using the formula: SBR = 20 × log_10_(*S*/*B*).

### Animal preparation

#### Animals

All animal experiments were conducted in accordance with the protocol approved by the Institutional Animal Care and Use Committee (IACUC) of the Department of Laboratory Animal Science at Fudan University (Approval No. 2022JS-ITBR-013).

Unless otherwise specified, this study utilized male wild-type C57BL/6JNifdc mice (Stock No. 219, Charles River Laboratories, China) and CaMKII-Cre:Ai162 transgenic mice, which express GCaMP6s in excitatory neurons. The CaMKII-Cre:Ai162 mice were generated by in-house breeding, crossing CaMKII-Cre mice (Stock No. C001015, Cyagen Biosciences) with Ai162 mice (Stock No. 031562, The Jackson Laboratory).

#### Viral injection

For wild-type mice, viral injections were performed at 7-8 weeks of age. Mice were anesthetized with 1.5% isoflurane (R510-22-10, RWD Life Science) in oxygen and secured on a heating pad in a stereotaxic apparatus (68803, RWD Life Science). Erythromycin ophthalmic ointment was applied to protect the eyes. The scalp was incised to expose the skull, which was then leveled in the mediolateral and anteroposterior planes using bregma and lambda as reference points. A 0.5 mm craniotomy was made at the target coordinates using a dental drill (78001 Microdrill, RWD Life Science). The virus was injected using a pulled glass micropipette connected to a 10 µL microsyringe (701N, Hamilton), controlled by a microinjection pump (KDS 130, KD Scientific). The virus was delivered at a rate of 50 nL/min, and the pipette was left in place for 10 min post-injection to prevent backflow. Mice were kept under continuous anesthesia throughout the procedure. Following the injection, the scalp was sutured.

For astrocyte imaging, the injection site was targeted to 1.5 mm posterior to bregma and 2 mm lateral to the midline (right hemisphere). The injected virus was AAV-GfaABC1D-GCaMP6s-P2A-tdTomato-WPREs (titer: 5 × 10^12^ vg/mL, diluted 1:1; PT-4766, BrainVTA). The virus was injected at one depth from the dural surface: 300 nL at 0.3 mm.

For neuronal imaging in the motor cortex, the injection site was targeted to 1.66 mm anterior to bregma and 0.4 mm lateral to the midline (right hemisphere). The injected virus was pAAV-hSyn-GCaMP6s WPRE (titer: 3 × 10^13^ vg/mL, diluted 1:3; Cat. No. E1273, Obiosh). The virus was injected at three depths from the dural surface: 100 nL at 0.9 mm, and 300 nL at 1.4 mm.

For neuronal imaging in the somatosensory cortex, CaMKII-Cre mice were used. The injection site was targeted to 0.38 mm anterior to bregma and 1.8 mm lateral to the midline (right hemisphere). The injected virus was pAAV-hSyn-FLEX-jGCaMP8s-WPRE (titer: 1 × 10^12^ vg/mL, diluted 1:1; Cat. No. E1273, Obiosh). The virus was injected at three depths from the dural surface: 300 nL at 0.5 mm, 300 nL at 1.1 mm and 300 nL at 1.5 mm.

For neuronal imaging in the PFC, the injection site was targeted to 1.5 mm anterior to bregma and 0.25 mm lateral to the midline (left hemisphere). The injected virus was pAAV-hSyn-GCaMP6s (titer: 3 × 10¹³ vg/mL, diluted 1:3; Cat. No. E1273, Obiosh). The virus was injected at three depths from the dural surface: 100 nL at 0.5 mm, 200 nL at 1.0 mm, and 300 nL at 1.5 mm.

#### Cranial window implantation

Similar to the viral injection procedure, mice were anesthetized and secured in the stereotaxic apparatus. The scalp was resected, and the skull was leveled. A circular craniotomy was performed using a dental drill, leaving the dura mater intact. Bleeding was controlled with an absorbable hemostatic agent. A custom-made glass window, consisting of a coverslip (150 µm thick) bonded to a glass ring (150 µm thick), was placed over the exposed brain area and sealed with tissue adhesive (Vetbond, 3M). Finally, a custom-made titanium headplate was attached to the skull using cyanoacrylate instant adhesive (5800, ergo, Switzerland). Dental cement (430205, New Century Dental Material Co., Ltd, Shanghai, China) was used for additional fixation of the headplate.

For transgenic mice designated for calcium imaging and wild-type mice designated for vasculature labeling, cranial window implantation was performed at 8-10 weeks of age. A 7 mm craniotomy was created, and a glass window, consisting of a 6.5 mm coverslip bonded to a glass ring (7.5 mm outer diameter, 6.0 mm inner diameter), was implanted.

For wild-type mice that received viral injections, the chronic cranial window was implanted 21 days after viral injection. A 4 mm craniotomy, centered over the injection site, was performed. A glass window, composed of a 3.5 mm coverslip bonded to a glass ring (5.5 mm outer diameter, 3.5 mm inner diameter), was then implanted.

After the surgical procedure, mice were housed for a seven-day recovery period. For post-operative care, dexamethasone (ST1254, Beyotime, Shanghai, China) and cefazolin (C832409, Macklin, Shanghai, China) were administered daily via intraperitoneal injection at a dose of 5 mg/kg body weight. Stock solutions were prepared at 0.5 mg/mL in sterile saline.

#### Vascular fluorescent labeling

To label the vasculature, wild type mice that had previously undergone a 7 mm craniotomy were anesthetized with isoflurane and subsequently received a retro orbital injection of 200 µL of 5% fluorescein isothiocyanate conjugated dextran (FITC Dextran; FD70S, Sigma Aldrich) dissolved in physiological saline. Imaging through the cranial window was completed within two hours post injection.

### Image processing and analysis

#### Image denoising

Unless otherwise specified, all images were processed with the OptiCal model to remove periodic noise. Any removal of random noise is specifically noted.

#### Volume rendering of cerebrovasculature

Volumetric imaging of mouse cerebrovasculature was performed with a z-axis step size of 5 µm. At each depth layer, 10 frames (Extended Data Fig. 2a) or 30 frames (Fig. 3a) were acquired and averaged following rigid-body registration. The depth stack was aligned using a custom MATLAB script to correct for axial drift, then imported into Imaris software (v10.1.0, Oxford Instruments) for volumetric reconstruction and maximum-intensity projection visualization.

#### Blood flow velocity measurement

Capillary blood flow was monitored using line-scan imaging at a sampling rate of 1,000 Hz. The acquired data were processed to generate kymographs, and blood flow velocity was calculated using a Fourier transform-based algorithm (Extended Data Fig. 6).

#### Volume rendering of GCaMP6s-labeled neurons

For Extended Data Fig. 7d, volumetric imaging of GCaMP6s-labeled neurons was performed from 0 to 965 µm depth with a z-axis step size of 5 µm. The depth stack was imported into Imaris software for volumetric reconstruction.

For Fig. 5b, both conventional and FPA imaging were performed from 0 to 1,700 µm depth with a z-axis step size of 5 µm. For conventional imaging (left), 20 frames were acquired at each depth layer and averaged following rigid-body registration. For FPA imaging (right), 200 frames were acquired at each depth layer and processed to generate 20 reconstructed frames. The depth stacks from both imaging modes were processed using a custom MATLAB script to generate X-Z maximum-intensity projections for side-view visualization.

#### Calcium signal extraction

Cell segmentation, signal extraction, and Δ*F*/*F*_0_ calculations were performed using custom software developed for our ROI imaging module. Neuronal ROIs were automatically identified from average intensity projections, standard deviation projections, and maximum-intensity projections of the calcium imaging movies, followed by manual verification and correction. For signal extraction, the fluorescence intensity of each ROI was calculated by averaging the top 30% of brightest pixels within the ROI mask for every frame. To track the same neurons across multiple imaging sessions, the same ROI masks were applied to align the data.

Then Δ*F*/*F*_0_ was computed using the following formula:

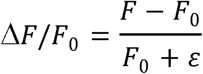

where *F* represents the instantaneous fluorescence intensity, and the baseline fluorescence (*F*_0_) is determined by averaging the fluorescence values between the 5th and 15th percentiles of the entire signal trace. A small constant (ε = 2.2204 × 10¹) was added to the denominator to prevent division by zero.

#### Spike inference

We inferred spikes from the fluorescence traces using the OASIS algorithm^35^. The algorithm employed an autoregressive model of order 1 (AR1) with a decay time constant of 1.5 s. The minimum spike magnitude threshold was determined automatically based on the estimated noise level of each fluorescence trace. This procedure yielded the denoised calcium traces and the inferred spike train. Calcium transients were identified from the denoised traces using MATLAB’s findpeaks function for quantitative analysis.

#### Photon count estimation

To estimate the number of fluorescence photons collected in our large-FOV 3PM, we first calibrated the detection system such that a single detected photon corresponded to an average grayscale value of 338. This conversion factor was derived from single-photon response calibration using fluorescein. We then converted the extracted raw fluorescence intensity values to photon counts by dividing them by this conversion factor.

#### Calculation of SNR

To assess imaging quality, we calculated SNR for three types of measurements: cerebrovasculature structure, calcium signal structure, and neuronal calcium dynamics.

For vascular structure imaging, we manually placed line ROIs perpendicular to blood vessels using ImageJ software (v2.14.0/1.54p, National Institutes of Health). Each line ROI had a fixed length of 60 pixels (≈ 33 μm) and was drawn across the vessel axis. For calcium signal structure imaging, line ROIs with the same dimensions (60 pixels ≈ 33 μm) were similarly drawn across neuronal structures. For calcium dynamics, SNR was calculated directly from the fluorescence traces of individual neurons extracted by our analysis pipeline.

The SNR calculation method was identical across all three measurements. The signal amplitude (*S*) was defined as the difference between the 95th percentile and the median fluorescence intensity of each trace. The noise level (*N*) was estimated from the median absolute difference between adjacent time points:

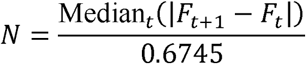

where division by 0.6745 converts the median absolute deviation to an equivalent standard deviation for normally distributed noise. Only positive differences were included in the calculation. The SNR was computed in decibels as:

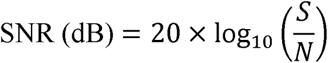

#### Heat-induced tissue damage assessment

Mice were subjected to continuous imaging for 20 min using either conventional imaging (120 mW) or ROI imaging (12 mW). Twenty to twenty-four hours later, the mice were transcardially perfused with ice-cold saline, followed by 4% paraformaldehyde (PFA). The brains were rapidly dissected and post-fixed in 4% PFA at 4 °C overnight. After cryoprotection in 30% sucrose solution for at least 24 h, the brains were embedded in optimal cutting temperature (OCT) compound (4583, Sakura Finetek) and sectioned into 25 μm-thick coronal slices using a cryostat (CM1950, Leica).

For immunohistochemistry, brain sections were washed in phosphate buffered saline (PBS) for 5 min and then blocked for 1 h at room temperature in blocking solution containing 5% normal donkey serum and 3% Triton X 100 in PBS. Sections were incubated overnight at 4 °C with the appropriate primary antibodies diluted in the blocking solution. After primary incubation, sections were washed and then incubated for 2 h at room temperature with species appropriate, fluorophore conjugated secondary antibodies diluted in blocking solution. Following washes with PBS (3 × 10 min), sections were mounted onto glass slides and coverslipped with an anti fade mounting medium containing DAPI (P0131, Beyotime Biotechnology). Sections were imaged using a slide scanner (Slideview VS200, Olympus). To check the expression of GCaMP6s, primary rabbit anti-GFP antibody (1:500, A-11122, Invitrogen) and secondary Alexa Fluor 488-conjugated donkey anti-rabbit (1:500, 711-545-152, Jackson) were used. To check the expression of HSP70, primary mouse anti-HSP70/72 antibody (1:400, ADI-SPA-810-D, Enzo Life Science) and secondary Alexa Fluor 594-conjugated donkey anti-mouse (1:500, 715-585-151, Jackson) were used.

Photodamage was quantified by comparing the fluorescence intensity of HSP70/72 immunostaining between the laser-exposed ipsilateral hemisphere and the non-exposed contralateral hemisphere, which served as an internal control. Specifically, 0.5 × 0.5 mm^2^ ROIs were defined in the corresponding cortical areas of both hemispheres using ImageJ software. The extent of damage was expressed as the relative change in HSP70/72 fluorescence intensity (Δ*I* / *I*_0_), calculated as: Δ*I* / *I*_0_ = (*I* − *I*_0_) / *I*_0_, where *I* is the mean intensity of the ipsilateral ROI and *I*_0_ is the mean intensity of the contralateral ROI.

### *In vivo* functional imaging in PFC

#### Behavioral paradigm

Mice were head-fixed within a custom-built apparatus. Each experimental session began with a 5-min baseline recording, followed by a 10-min period of intermittent air puff stimulation. During the stimulation period, each trial consisted of a 15-s air puff stimulus followed by a 45-s inter-stimulus interval (ISI). Throughout the session, the body movements of the mice were monitored using an infrared camera operating at 30 frames per second.

#### Neural population analysis

Neural population dynamics were downsampled using PCA. To investigate the relationship between population activity and behavior, we computed the Spearman correlation coefficients among three variables: (1) the global average activity, (2) the first principal component (PC1), and (3) motion energy.

Motion energy was quantified based on the temporal dynamics of frame-to-frame image correlations. For each pair of consecutive frames, we computed the Spearman correlation coefficient (*r*). We then converted this to a decorrelation measure (1 − *r*), applied z-score normalization across the time series, and took the absolute value to obtain the final motion energy metric.

### Statistical analysis and quantification

In all box plots, center lines represent medians, box boundaries represent upper and lower quartiles, and whiskers extend to 1.5 times the interquartile range.

All data processing and statistical analyses were performed using custom-written scripts in MATLAB (R2023b, Mathworks). Normally distributed continuous data are presented as mean ± SEM. All statistical details (the size and type of individual samples, n) are specified in figure legends.

## Data availability

All data generated during this study are included in the published article, and raw data are available from the corresponding author upon request. Any additional information required to reanalyze the data reported in this paper is available from the lead contact upon request.

## Code availability

The ROI imaging module software and OptiCal denoising algorithm are available at: https://github.com/Achuan-2/3PM_ROI_imaging_module.

## Acknowledgements

This work was supported by the National Natural Science Foundation of China (32471142, T2222006, 32541017), Science and Technology Commission of Shanghai Municipality (22JC1403100).

## Author Contributions

B.L. supervised the project and co-designed the study with J.X.S., S.P.L., and S.S.Y. B.L., S.P.L. and M.Z. designed and built the imaging system. J.X.S. developed the ROI imaging module software. S.P.L. developed the OptiCal denoising model. J.X.S., S.S.Y., Y.F.Z., X.Y.G., Y.G.Z. and C.Y.L performed the animal surgeries. J.X.S. and S.P.L. analyzed the data. J.X.S., S.P.L., and B.L. wrote the manuscript with input from all authors.

## Extended data

**Extended Data Fig. 1.**
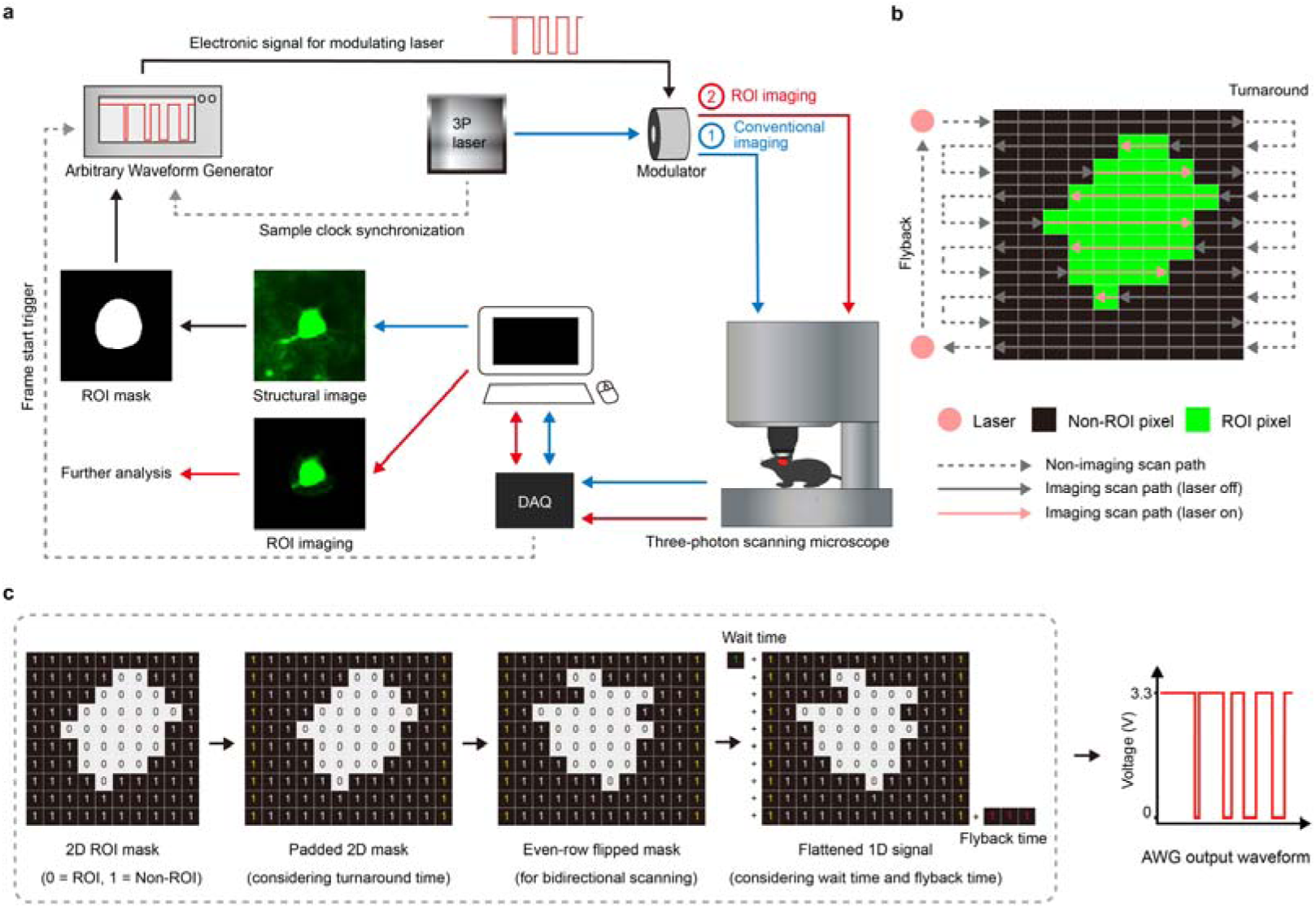
Hardware architecture and signal generation principle of the ROI imaging system. **a,** Hardware architecture and data flow of the ROI imaging module. A structural reference image is first acquired through conventional imaging or FPA imaging (blue path ①). A 2D binary ROI mask is generated from this image after cell segmentation and then converted into a 1D modulation signal loaded into the AWG. The AWG output drives a laser modulator (e.g., AOM) to enable high-speed laser gating according to preset ROI positions (red path ②). For precise pixel-level synchronization, the AWG sampling clock is synchronized with the laser’s master clock, and the galvanometer’s frame-start trigger initiates waveform output from the AWG. **b,** Bidirectional raster scanning path and laser modulation for ROI imaging. The schematic shows the typical bidirectional scanning path, including imaging time and non-imaging time (turnaround and flyback with laser off, dashed arrows). Green pixels indicate the ROI where the laser is active; black pixels indicate non-ROI areas where the laser is blanked. Laser is switched on only when the focus is inside the ROI (pink path). **c,** Generation of the 1D temporal modulation waveform from the 2D ROI mask. The 2D ROI mask (0 = ROI, 1 = Non-ROI) is first padded with additional pixels to account for turnaround time. For bidirectional scanning, even-numbered rows are horizontally flipped to match the reversed scan direction. The processed 2D mask is then flattened row by row into a 1D digital signal, with additional signals added for wait time and flyback time. This 1D signal is loaded into the AWG memory and converted into a voltage waveform. Low voltage (0 V) enables laser transmission at ROI pixels, whereas high voltage (3.3 V) blocks laser transmission at non-ROI pixels and non-imaging time.

**Extended Data Fig. 2.**
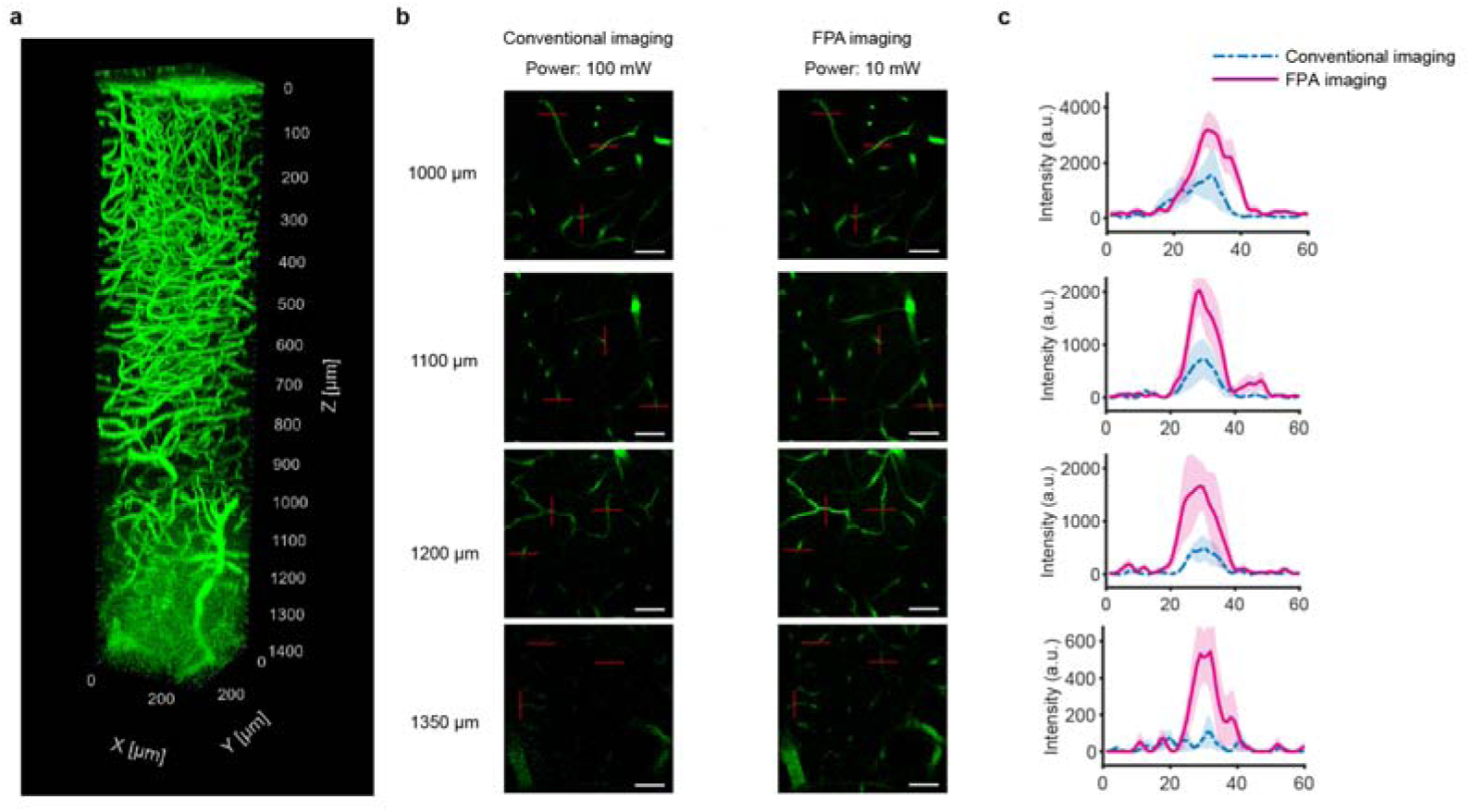
FPA imaging improves deep vascular imaging quality with lower laser power. **a,** Structural imaging of mouse cerebrovasculature. The 3D vascular structure was recorded with a FOV of 285 × 285 μm^2^, a Z-step size of 5 µm, and an imaging depth ranging from 0 to 1,400 μm. Each depth layer was averaged from 10 frames. **b,** Comparison between conventional and FPA imaging. Representative images acquired using conventional imaging (100 mW, left) and FPA imaging (12 mW, right) at depths from 1,000 µm to 1,350 µm. Each image represents a maximum-intensity projection of 10 frames. Scale bar, 50 μm. **c,** Signal intensity profile. Fluorescence intensity was measured along the red line shown in **b**. Lines represent mean signal intensity, and shaded areas indicate SEM.

**Extended Data Fig. 3.**
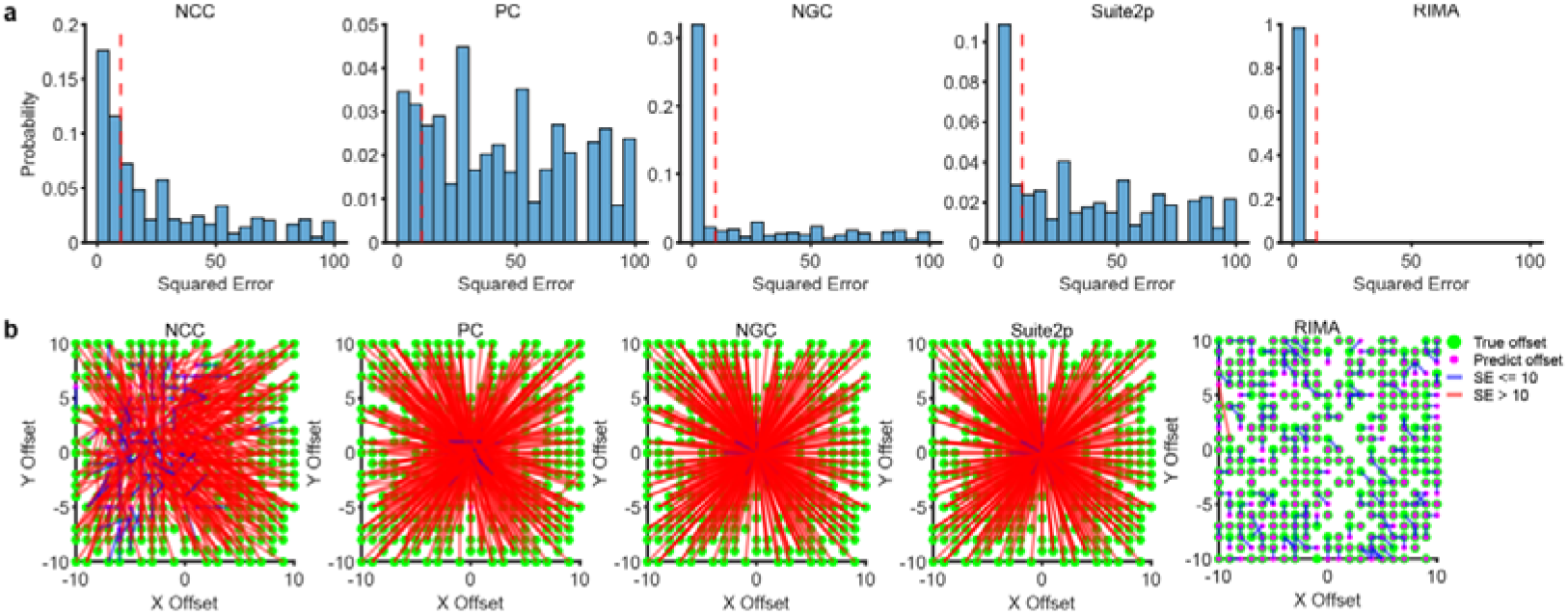
Quantitative comparison of registration accuracy of different algorithms on conventional and ROI imaging data. **a,** Histogram of squared registration errors of five registration algorithms on ROI imaging data. **b,** Visualization of registration results from the five algorithms in **a**. All experiments were performed on *n* = 50 calcium imaging videos with applied displacements.

**Extended Data Fig. 4.**
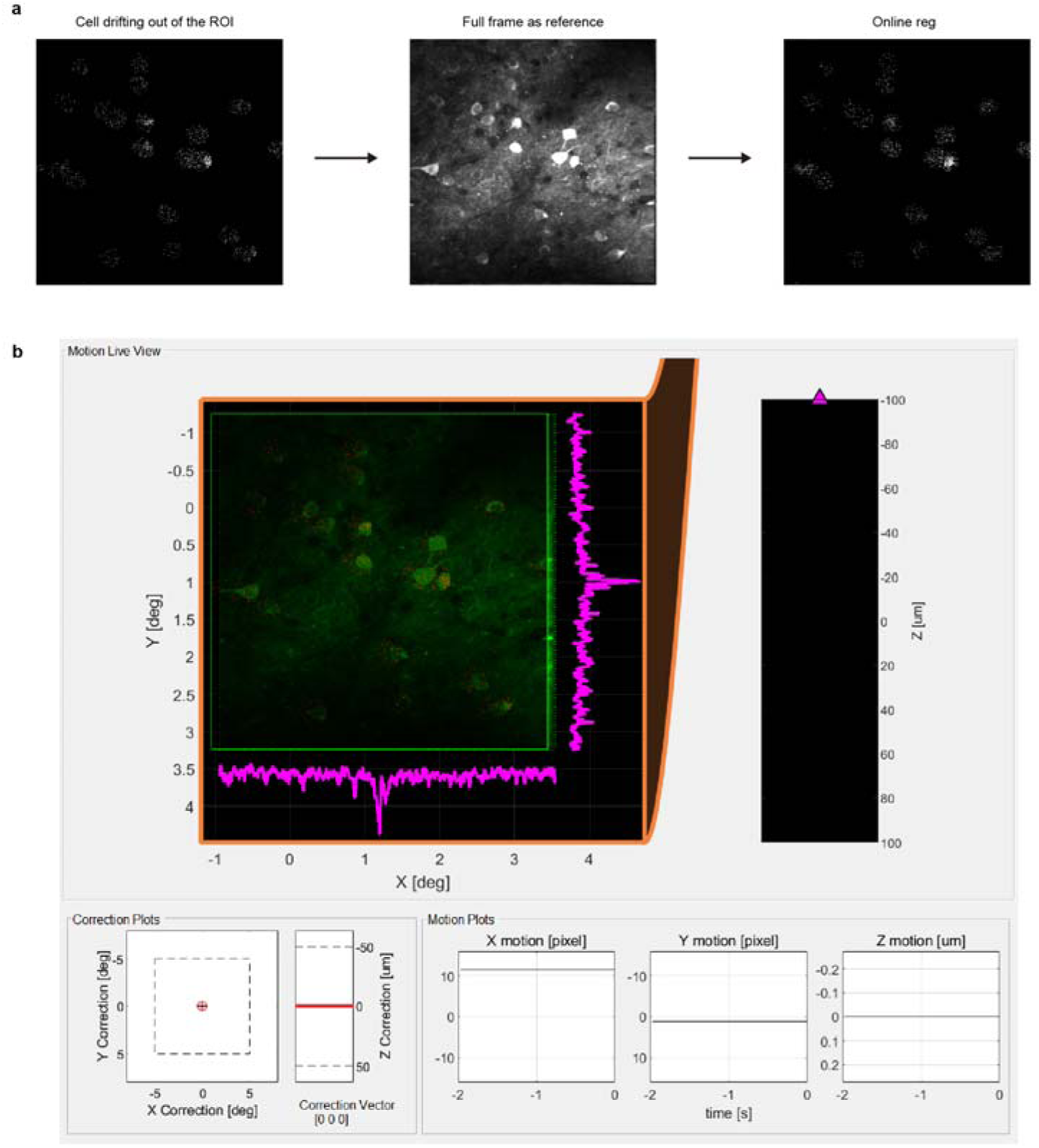
Online registration for ROI imaging. a, Schematic of the Online-Reg working principle. Left, cells begin to deviate from the preset ROI mask due to animal movement during ROI imaging. Middle, a pre-acquired full-field image from FPA imaging is used as a stable registration reference. Right, the Online-Reg algorithm corrects scanning coordinates in real time to keep the ROI continuously locked on target cells. **b,** GUI for real-time motion correction. The main view displays registration status in real time, where the green channel represents the reference image and the red channel represents the currently acquired ROI signal; magenta curves at the edges show signal projections along X and Y axes.

**Extended Data Fig. 5.**
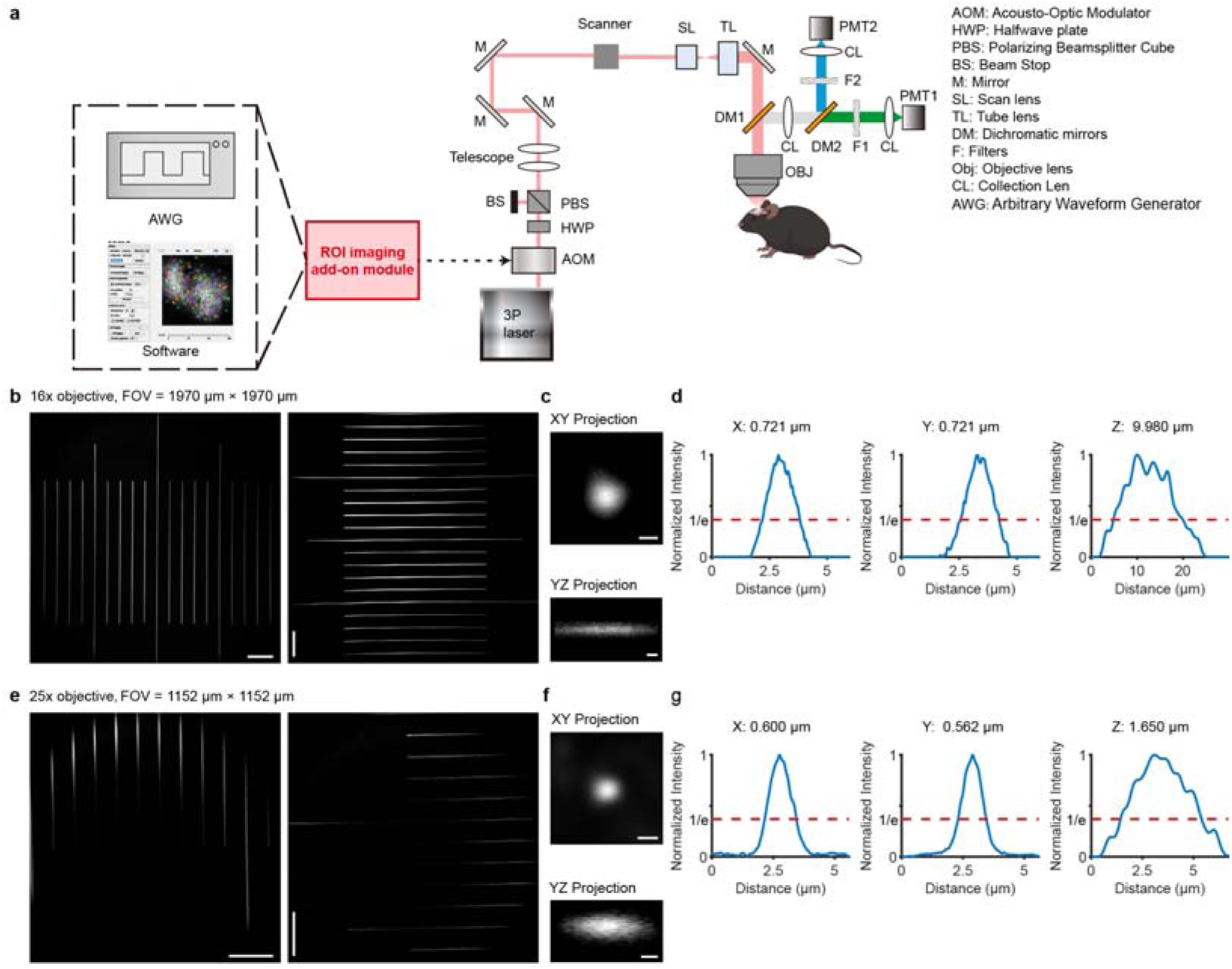
Design and performance demonstration of the customized 3PM. **a,** Optical path schematic of the customized large-FOV three-photon microscope. **b-d,** System performance characterization using a 16× water-immersion objective (NA = 0.8, 3 mm working distance; CFI75 LWD 16X W, Nikon). (**b)** The FOV measured with a standard calibration slide is 1,970 × 1,970 μm^2^. **(c)** XY and YZ plane projections of single fluorescent microsphere imaging for PSF measurement. **(d)** Normalized intensity profiles of the microsphere shown in **c** along the X, Y, and Z axes. The system lateral resolutions are 0.72 µm (X) and 0.72 µm (Y), and the axial resolution (Z) is 9.98 µm (defined as the 1/e width of the normalized intensity profile). **e-g,** System performance characterization using a 25× water-immersion objective (NA = 1.05, 2 mm working distance; XLPLN25XWMP2, Olympus). **(e)** The FOV measured with a standard calibration slide is 1,152 × 1,152 μm^2^. **(f)** XY and YZ plane projections of single fluorescent microsphere imaging for PSF measurement. **(g)** Normalized intensity profiles of the microsphere shown in **f** along the X, Y, and Z axes. The system lateral resolutions are 0.6 µm (X) and 0.56 µm (Y), and the axial resolution (Z) is 1.65 µm (defined as the 1/e width of the normalized intensity profile). Scale bars, 200 μm in **b**,**e**; 1 μm in **c**,**f**.

**Extended Data Fig. 6.**
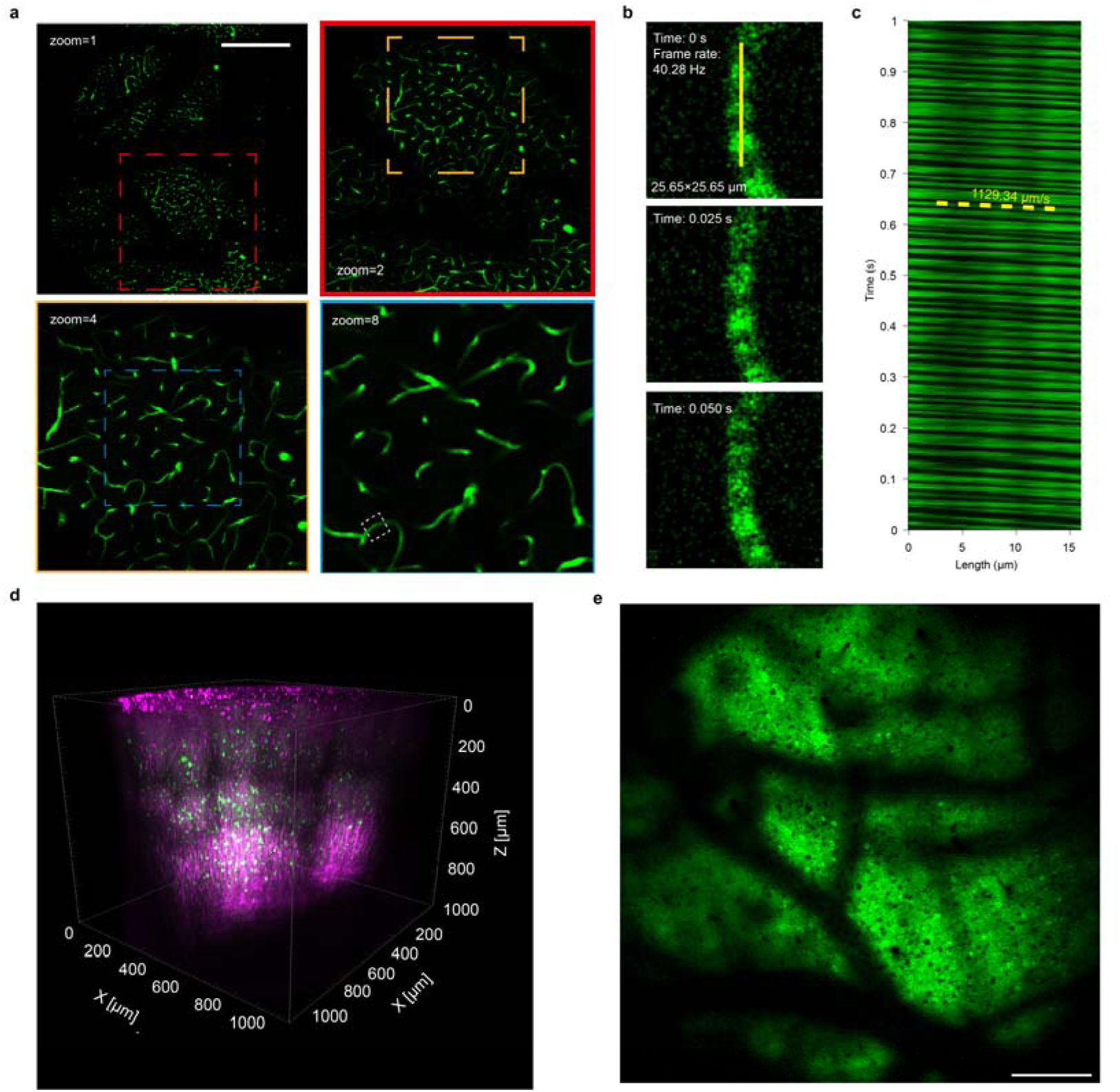
*In vivo* imaging of mouse cortical vasculature and blood flow velocity measurement. **a**, Cortical vasculature imaging at multiple zoom levels. Representative images acquired at a depth of 300 µm using four zoom settings. The FOV decreases from 1,970 × 1,970 µm² (zoom = 1, top left) to 246 × 246 µm² (zoom = 8, bottom right). Scale bar, 500 µm. **b**, Time-lapse imaging of a single capillary acquired in ROI scanning mode at 40.28 Hz. Unlabeled red blood cells appear as dark shadows flowing through the fluorescent plasma. Three consecutive frames are shown at the indicated time points. **c**, Kymograph of capillary blood flow velocity generated from a 1-kHz line scan along the central axis of the capillary (yellow line in **b**). The diagonal streaks represent trajectories of individual RBCs, and their slopes correspond to a flow velocity of 1129.34 μm/s. **d**, 3D reconstruction of deep cortical structure in a CaMKII-Cre:Ai162 transgenic mouse, covering a volume of 1,152 × 1,152 × 965 µm^3^ acquired with a 25× objective. The reconstruction visualizes third-harmonic generation signals (magenta) and fluorescence from GCaMP6s-expressing neurons (green). **e,** A representative average image of GCaMP6s-expressing neurons at 200 µm depth in the mouse cerebral cortex. Scale bar, 200 µm.

**Extended Data Fig. 7.**
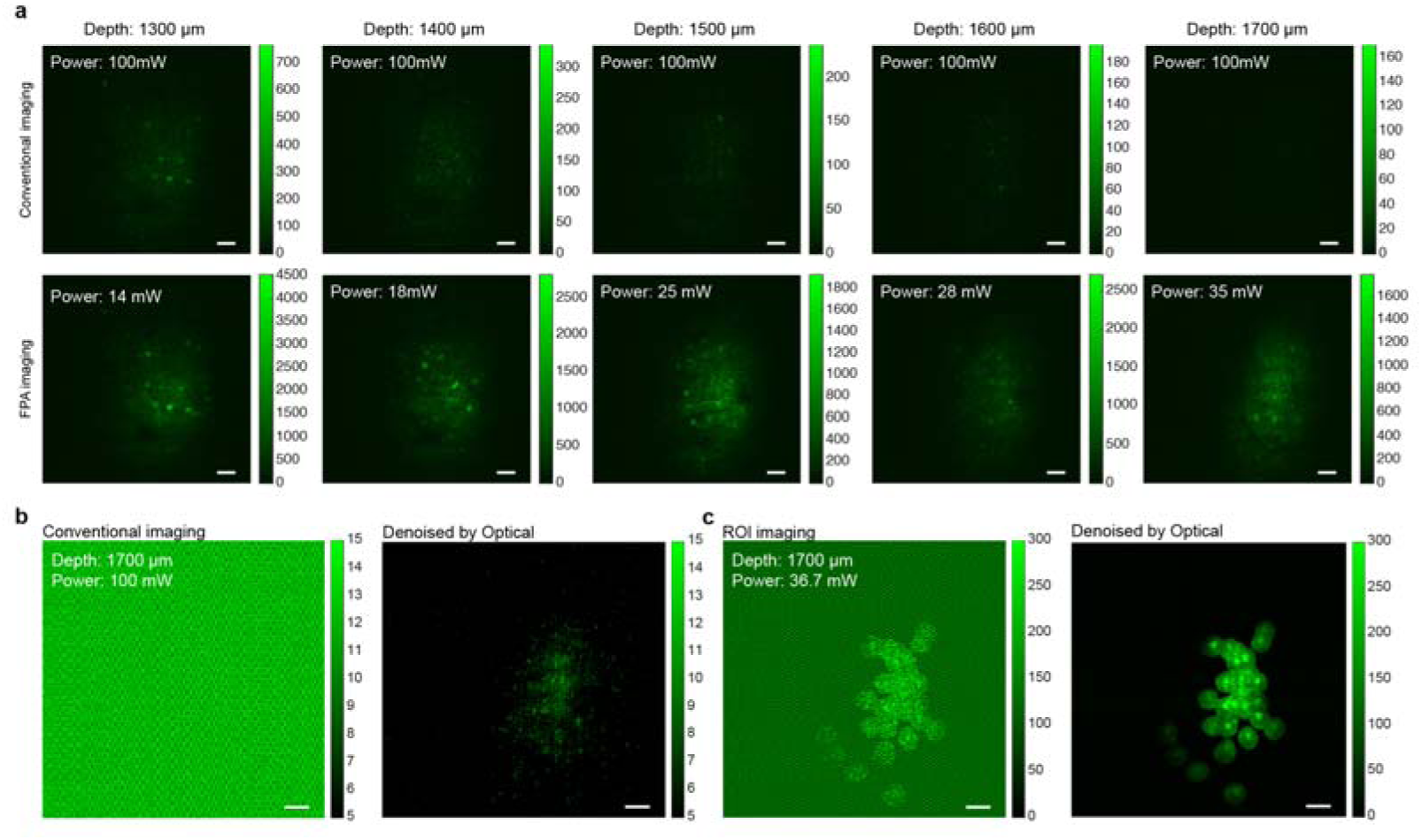
Comparison of deep imaging performance and denoising efficacy. **a**, Representative projection images comparing conventional imaging (top) with FPA imaging (bottom) at various depths in the deep PFC cortex, from 1,300 µm to 1,700 µm. Scale bar, 50 μm. **b**, Representative frame from conventional imaging at 1,700 µm. Left, raw image. Right, image after removal of both periodic and random noise using OptiCal. Scale bar, 50 μm. **c**, Representative frame from ROI imaging at 1,700 µm. Left, raw image. Right, image after removal of both periodic and random noise using OptiCal. Scale bar, 50 μm.

