## Supplementary Note for "A plug-and-play ROI imaging module and deep-learning denoising framework extend three-photon microscopy to 1.7 mm depth"

### Supplementary Note 1 | Calculation of average power in ROI imaging

In ROI imaging, an acousto-optic modulator (AOM) gates the excitation laser. The laser is active only during pixel dwell times within predefined ROIs and is turned off during all other periods. These inactive periods include scanning of non-ROI regions and scanner overhead times, such as line turnarounds and frame flybacks.

Therefore, the effective average laser power delivered to the sample during ROI imaging ($P_{\text{ROI}}$) can be expressed as:

$$P_{\text{ROI}}=P_{\text{conv}}\cdot F_{\text{fill}}\cdot R_{\text{ROI}}$$

where $P_{\text{conv}}$ is the average power for conventional full-field imaging (measured with a power meter placed after the objective lens), $R_{\text{ROI}}$ is the fraction of the total FOV area occupied by the ROIs, and $F_{\text{fill}}$ is the scanner fill factor.

The fill factor, $F_{\text{fill}}$, represents the ratio of the active pixel acquisition time to the total time required for a full-frame scan. For a standard bidirectional raster scanner, $F_{\text{fill}}$ is determined by the specific scanning parameters as follows:

$$F_{\text{fill}}=\frac{H\cdot W}{t_{\text{wait}}+H\cdot\left( W+t_{\text{turn}} \right)+t_{\text{fly}}}$$

where:

- $H$ is the image height in pixels.
- $W$ is the image width in pixels.

- $t_{\text{wait}}$ is the initial wait time before scanning begins.

- $t_{\text{turn}}$ is the turnaround time for the bidirectional scanner to reverse direction at the end of each line.

- $t_{\text{fly}}$ is the flyback time required for the scanner to return from the end of a frame to the start of the next one.

All time parameters ($t_{\text{wait}}$, $t_{\text{turn}}$, and $t_{\text{fly}}$) are measured in units of pixel dwell time. For the scanning parameters used in our system, we calculated $F_{\text{fill}}$ ≈ 0.9445.

### Supplementary Note 2 | Overcoming autocorrelation artifacts for ROI imaging data registration

Conventional calcium imaging registration algorithms, including normalized cross-correlation (NCC), phase correlation (PC), normalized gradient correlation (NGC), and Suite2p, perform poorly on ROI imaging data. These methods fail because ROI images contain large, stationary non-illuminated regions alongside small illuminated cellular structures subject to motion displacement. The fixed mask regions generate strong artifacts that often overwhelm the cross-correlation signal from true cellular displacements, leading to registration failure (**Extended Data Fig. SN1**).

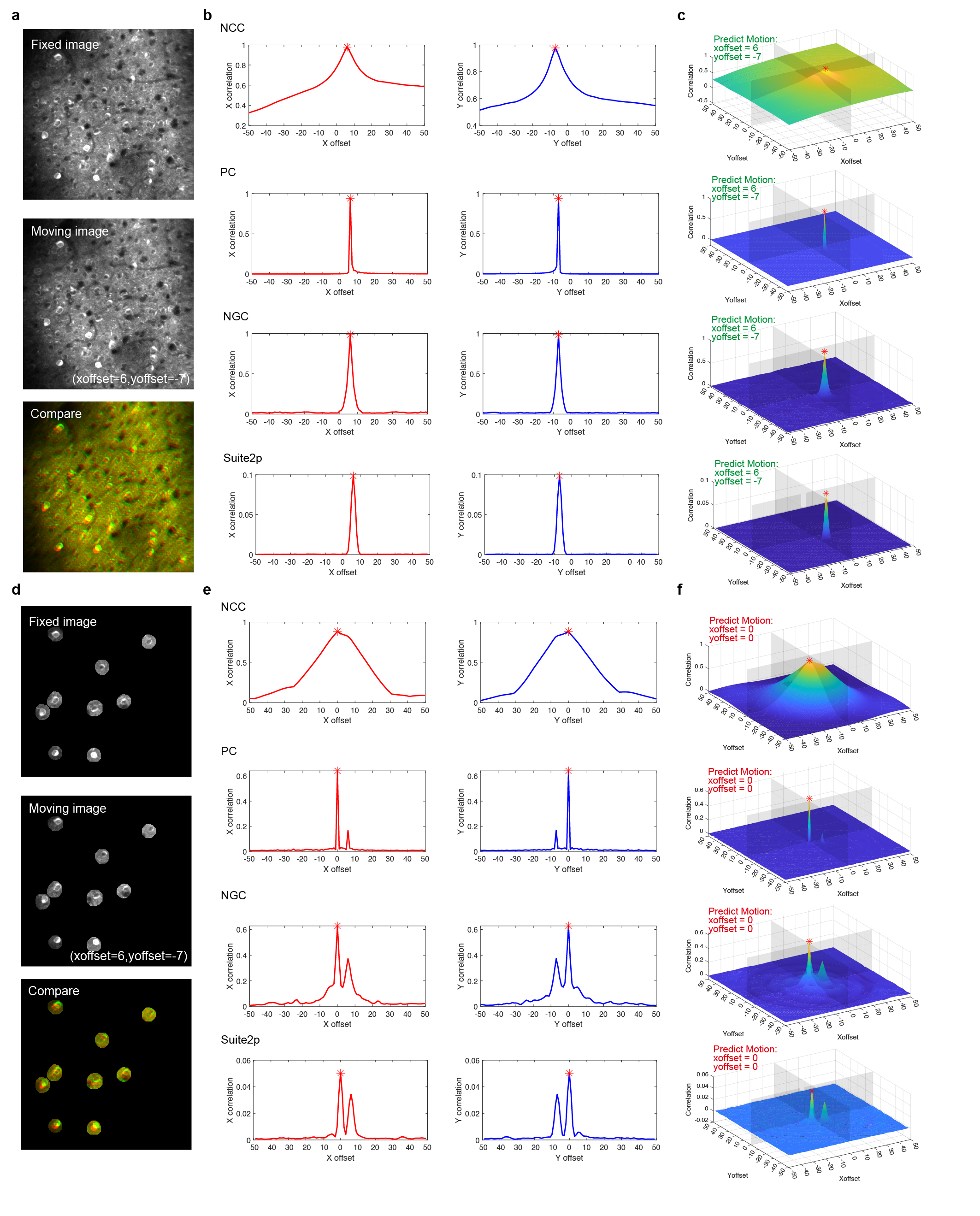

**Extended Data Fig. SN1 | Conventional registration algorithms fail on ROI imaging data. a,** Example full-field images: reference (top), moving (middle), and overlay after successful registration (bottom). **b,c,** 1D profiles (**b**) and 2D surfaces (**c**) of correlation maps for four conventional algorithms. All methods accurately identify the motion displacement in full-field data. **d,** ROI-masked images derived from (**a**), with the same displacement. **e,f,** 1D profiles (**e**) and 2D surfaces (**f**) of correlation maps for the ROI images. The artifact causes conventional algorithms to fail by incorrectly identifying zero displacement. Red asterisks indicate the peak position, which represents the predicted motion offset.

To address this challenge, we developed RIMA, a registration algorithm tailored for ROI imaging data (see Methods for implementation details). RIMA uses a high-SNR, full-field structural image obtained via FPA imaging as the reference template to align the subsequent ROI frames. This approach effectively suppresses artifacts and enables accurate motion estimation, achieving a registration accuracy of 99.5 ± 0.3% (**Fig. 1f**, **Extended Data Fig. SN2**). RIMA was therefore adopted as the dedicated algorithm for both online and offline registration.

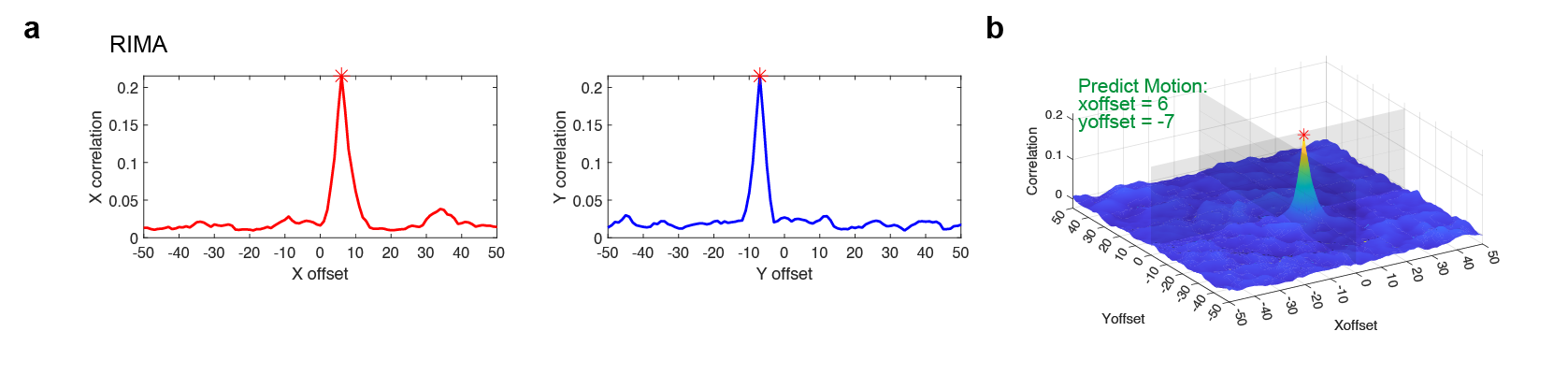

**Extended Data Fig. SN2 | RIMA overcomes artifacts for accurate ROI registration. a,b,** 1D profile (**a**) and 2D surface (**b**) of the correlation map generated by RIMA for the ROI-masked images shown in Extended Data Fig. SN1d. RIMA suppresses the mask-induced artifact, producing a single, clear cross-correlation peak that accurately identifies the true motion displacement. Red asterisks indicate the peak position, which represents the predicted motion offset.
